# A One-Pot Biocatalytic Cascade to Access Diverse L-Phenylalanine Derivatives from Aldehydes or Carboxylic Acids

**DOI:** 10.1101/2024.12.06.627276

**Authors:** Shelby R. Anderson, Madan R. Gopal, Abigail P. Spangler, Michaela A. Jones, D’Jana R. Wyllis, Aditya M. Kunjapur

## Abstract

Non-standard amino acids (nsAAs) that are L-phenylalanine derivatives with aryl ring functionalization have long been harnessed in natural product synthesis, therapeutic peptide synthesis, and diverse applications of genetic code expansion. Yet, to date these chiral molecules have often been the products of poorly enantioselective and environmentally harsh organic synthesis routes. Here, we reveal the broad specificity of multiple natural pyridoxal 5’-phosphate (PLP)-dependent enzymes, specifically an L-threonine transaldolase, a phenylserine dehydratase, and an aminotransferase, towards substrates that contain aryl side chains with diverse substitutions. We exploit this tolerance to construct a one-pot biocatalytic cascade that achieves high-yield synthesis of 18 diverse L-phenylalanine derivatives from aldehydes under mild aqueous reaction conditions. We demonstrate addition of a carboxylic acid reductase module to this cascade to enable the biosynthesis of L-phenylalanine derivatives from carboxylic acids that may be less expensive or less reactive than the corresponding aldehydes. Finally, we investigate the scalability of the cascade by developing a lysate-based route for preparative-scale synthesis of 4-formyl-L-phenylalanine, a nsAA with a bio-orthogonal handle that is not readily market-accessible. Overall, this work offers an efficient, versatile, and scalable route with the potential to lower manufacturing cost and democratize synthesis for many valuable nsAAs.

## Introduction

Non-standard amino acids (nsAAs) are valuable building blocks for chemical diversification of products that are prepared by synthetic or biological means.^[1–3]^ Diverse commercially relevant applications have been enabled by the introduction of sidechain chemistries that do not belong to the standard twenty ribosomally translated amino acids. An illustrative though not exhaustive list includes the formation or augmentation of bioactive natural products^[4–6]^, synthesis of stimuli-responsive materials^[7,8]^, improved protein stability^[9]^, increased enzyme catalytic efficiency^[10]^, synthesis of artificial metalloenzymes^[11]^ and (photo)catalytic active sites^[12,13]^, design of metal-responsive protein switches^[14]^, immunochemical termination of protein self-tolerance^[15]^, bio-orthogonal conjugation of therapeutics^[16–18]^, and intrinsic biological containment of genetically modified organisms^[19–22]^. Notably, each of the applications described above has prominently featured derivatives of L-phenylalanine, which are the most common class of nsAAs used in genetic code expansion approaches within live cells. However, efficient and cost-effective production of enantioenriched phenylalanine derivatives has been a longstanding challenge.

Biocatalysis presents a promising solution by utilizing enantioselective enzymes under mild reaction conditions, eliminating the need for protecting groups, and offering an increasingly scalable, sustainable, and economical approach compared to traditional catalytic routes.^[23]^ Two classes of PLP-dependent biocatalysts that exhibit wide substrate scope during synthesis of amino acids and that harness inexpensive precursors are L-threonine aldolases (L-TAs) and L-threonine transaldolases (L-TTAs). These catalyze aldol-like condensations of aldehydes and either L-glycine or L-threonine (L-Thr), respectively, to generate β-hydroxy α-amino acids (β-OH AAs). They exhibit activity on a broad range of aldehydes with aryl side chains containing diverse ring substitutions.^[24,25]^ L-TTAs are especially promising compared to L-TAs given their ability to form a glycyl quinonoid intermediate whose protonation is kinetically disfavored, enabling efficient desired reactivity with aryl aldehydes rather than undesired formation of glycine.^[26]^ Additionally, enzymatic oxidation or reduction of the acetaldehyde co-product of the L-TTA reaction offers a sink that can further shift equilibrium towards product formation independent of the chosen substrate.^[27]^ Through bioprospecting, we further discovered L-TTA homologs with improved function, such as high L-Thr affinity.^[24]^

While β-hydroxylated nsAAs could find substantial use in drug discovery, the formation of the β-hydroxy group results in chemical and steric differences from standard phenylalanine analogs and are currently less well-explored industrially. Therefore, we inquired about whether we could supply additional enzymes to catalyze the sequential removal of the β-hydroxy group contained within diverse phenylalanine derivatives that might be accessible via L-TTAs, ideally all in one pot. Due to their process intensification and sustainability, one-pot biocatalytic cascades are of high interest in the pharmaceutical and fine chemical industries^[28–32]^, having been used for biosynthesis of investigational treatments for HIV^[33]^ and cancer^[34]^ as well as for pharmaceutical synthons, particularly heterocyclic amines^[35,36]^. Additionally, numerous multi-enzymatic cascades have been developed to produce L- and D-α-amino acids, such as the industrially applied hydantoinase^[37]^ and acylase^[38]^, or broadly amidohydrolase, processes.^[32,39,40]^ Although these processes initialize from inexpensive precursors, they are typically limited by the narrow range of available substrates. Given this precedent, a one-pot biocatalytic cascade could be a solution with high industrial translatability for phenylalanine derivative manufacture if the biocatalysts exhibit broad tolerance and if the necessary precursors are cheap and readily available. However, for the reaction to proceed in the desired sequence without substantial byproduct formation, the cascade enzymes require a delicate balance of specificity; each enzyme in the cascade must exhibit sufficiently broad tolerance towards aryl ring substitution while maintaining selectivity for desired transformations at the α- and β-carbons. Additionally, the enzymes must exhibit limited crosstalk in the amino acid co-substrates that they harness for aldol condensation or for amine donation, respectively. Finally, for practicality each enzyme in the cascade must be sufficiently active as to require low micromolar enzyme concentrations throughout.

Here, we report on the remarkably broad substrate tolerance of two model natural PLP-dependent enzymes, a phenylalanine dehydratase and an aromatic amino acid aminotransferase, which we couple with a natural L-TTA to design a high-yield cascade capable of converting numerous non-native substrates to unnatural products of considerable demonstrated value. Despite previous work demonstrating a cascade which employs a TA and threonine deaminase, we study a phenylserine dehydratase to prevent consumption of the co-substrate L-threonine and overcome limitations of substrate scope.^[41,42]^ Thus, we study the phenylserine dehydratase from *Ralstonia pickettii* PS22 (*Rp*PSDH)^[43,44]^ coupled with an *E. coli* aromatic amino acid aminotransferase, TyrB (Scheme 1). The resulting one-pot cascade of purified enzymes generates phenylalanine derivatives with high yield and diversity from readily available and inexpensive aryl aldehyde precursors when coupled to an L-TTA from *Pseudomonas fluorescens* (ObiH). We further expand the versatility of our cascade through the addition of a promiscuous carboxylic acid reductase from *Segniliparus rotundus* (*Sr*CAR) for *in situ* generation of aryl aldehydes from carboxylic acids, which are often less expensive and more stable than aryl aldehydes. Finally, we show our system has potential for scale-up using clarified lysate from bacterial cells deficient in aldehyde reductases, achieving up to 64% yield of a poorly commercially accessible nsAA with a handle for bio-orthogonal conjugation, 4-formyl-L-phenylalanine, from 25 mM of the corresponding aldehyde precursor.

**Scheme 1.**
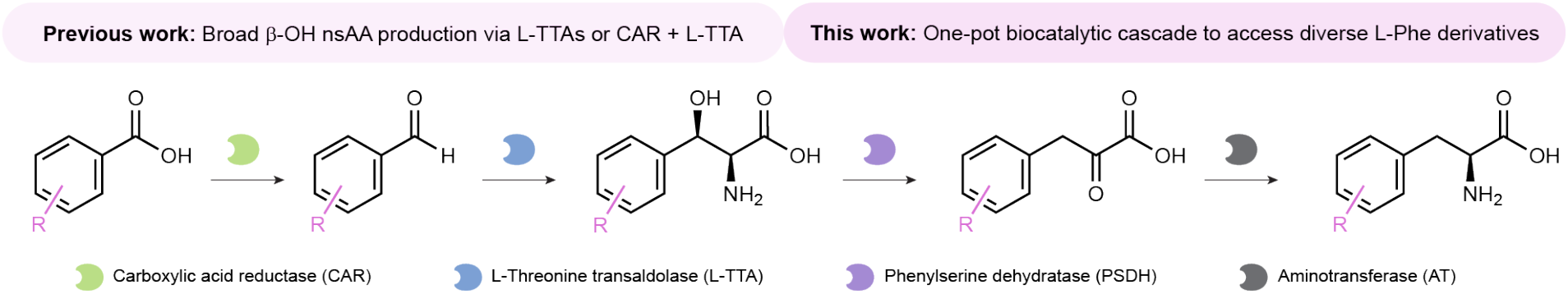
One-pot biocatalytic cascade to access diverse phenylalanine derivatives.

## Results and Discussion

### Identifying candidate enzymes for nsAA production

Towards identifying an enzyme capable of removing the β-hydroxy moiety from α-amino acids produced by an L-TTA catalyzed reaction, we considered the PLP-dependent phenylserine dehydratase from *Ralstonia pickettii* (*Rp*PSDH). While a relatively understudied enzyme in the literature, it has been shown to act on L-*threo*-phenylserine and was recently coupled to ObiH to form the associated keto acid from a β-OH leucine substrate.^[25,43,44]^ To evaluate *Rp*PSDH, we expressed a variant containing a N-terminal His-tag and purified it by nickel affinity chromatography for subsequent activity screening. We sought to confirm *Rp*PSDH activity on an aromatic β-hydroxylated α-amino acid in the presence of L-Thr, a co-substrate for the L-TTA reaction that is commonly supplied at superstoichiometric concentrations. Because few β-OH nsAAs are commercially available, we focused the initial screen on L-*threo*-phenylserine to inquire how this reaction might be affected by provision of L-Thr. We measured substrate depletion after addition of 2 μM *Rp*PSDH (0.07 mg/mL) to a reaction mixture containing 0.4 mM PLP and 1 mM L-*threo*-phenylserine with and without 100 mM L-Thr (Supporting Information, Figure S1). After two hours, we observed >95% of L-*threo*-phenylserine depletion under both conditions.

Given our ability to produce a prochiral α-keto acid after dehydration of the corresponding β-OH α-amino acid, we next turned to candidates for enantioselective amination at the α-carbon position. We hypothesized that a system in which we leveraged native aromatic amino transferases, should they exhibit broad specificity towards non-native substrates, might have greater relevance for scale-up. This hypothesis is supported by pathways such as the *de novo* biosynthesis of 4-nitrophenylalanine, which harness endogenous levels of *E. coli* aminotransferases.^[45]^ We evaluated the native *E. coli* aromatic amino transferase, TyrB, shown previously to proceed with excellent stereoselectivity at the α-carbon, including for non-standard L-phenylalanine derivatives, and is known not to accept L-threonine as an amine donor.^[46–48]^ After purification of N-terminal His-tagged TyrB, we confirmed its activity on phenylpyruvic acid and then compared its activity to one non-native ring-functionalized phenylpyruvic acid derivative, 4-nitrophenylpyruvic acid (**1d**). TyrB acted on both substrates rapidly, and we were excited to observe full conversion of supplemented 0.5 mM **1d** to the corresponding nsAA **1e** after 15 min in a reaction mixture with 10 mM L-glutamic acid (L-Glu) as the amine donor and 0.4 mM PLP (Supporting Information, Figure S2).

### One-pot nsAA production from aldehyde precursors

We next evaluated whether we could combine all three broadly specific PLP-dependent enzymes in one pot for a reaction cascade. Given our recent discovery of improved L-TTA expression with the addition of a SUMO-tag without compromised enzyme activity, we purified the L-TTA from *Pseudomonas fluorescens* with appended N-terminal Hisand SUMO tags, denoted s-ObiH.^[24]^ We conducted an initial test of reaction buffer compatibility and relative kinetics by supplying 2 mM of either phenylpyruvate, phenylserine, or benzaldehyde to a one-pot preparation of s-ObiH, *Rp*PSDH, and TyrB, confirming endpoint phenylalanine production in each condition (Supporting Information, Figure S3). Building upon this foundation, we investigated the ability of this candidate cascade to produce diverse L-phenylalanine derivatives from their associated benzaldehyde derivatives (Figure 1a). We varied the position, size, and electrostatic properties of aryl ring functional groups, ultimately selecting 18 commercially valuable or representative chemistries in order to ascertain the preferences of this candidate pathway. We added equimolar enzyme concentration (2 μM) of each purified enzyme (s-ObiH (0.13 mg/mL), *Rp*PSDH (0.07 mg/mL), and TyrB (0.09 mg/mL)) to a reaction mixture containing 0.4 mM PLP, 100 mM L-Thr, 25 mM L-Glu, and 2 mM aryl aldehyde precursor. We were excited to observe generally high yields with our reaction cascade after 12 h at 30 °C. Notably, seven out of the 18 chemistries exhibited yields exceeding 96%, with 13 chemistries (and thus the majority of chemistries tested) exhibiting over 75% yield without any optimization (Figure 1b). Further, the reactions proceeded with high enantioselectivity to produce L-phenylalanine derivatives, with greater than 99% enantiomeric excess (*e*.*e*.) for the 17 chemistries screened (**11e** was not analyzed) (Supporting Information, Figures S4-S21).

**Figure 1.**
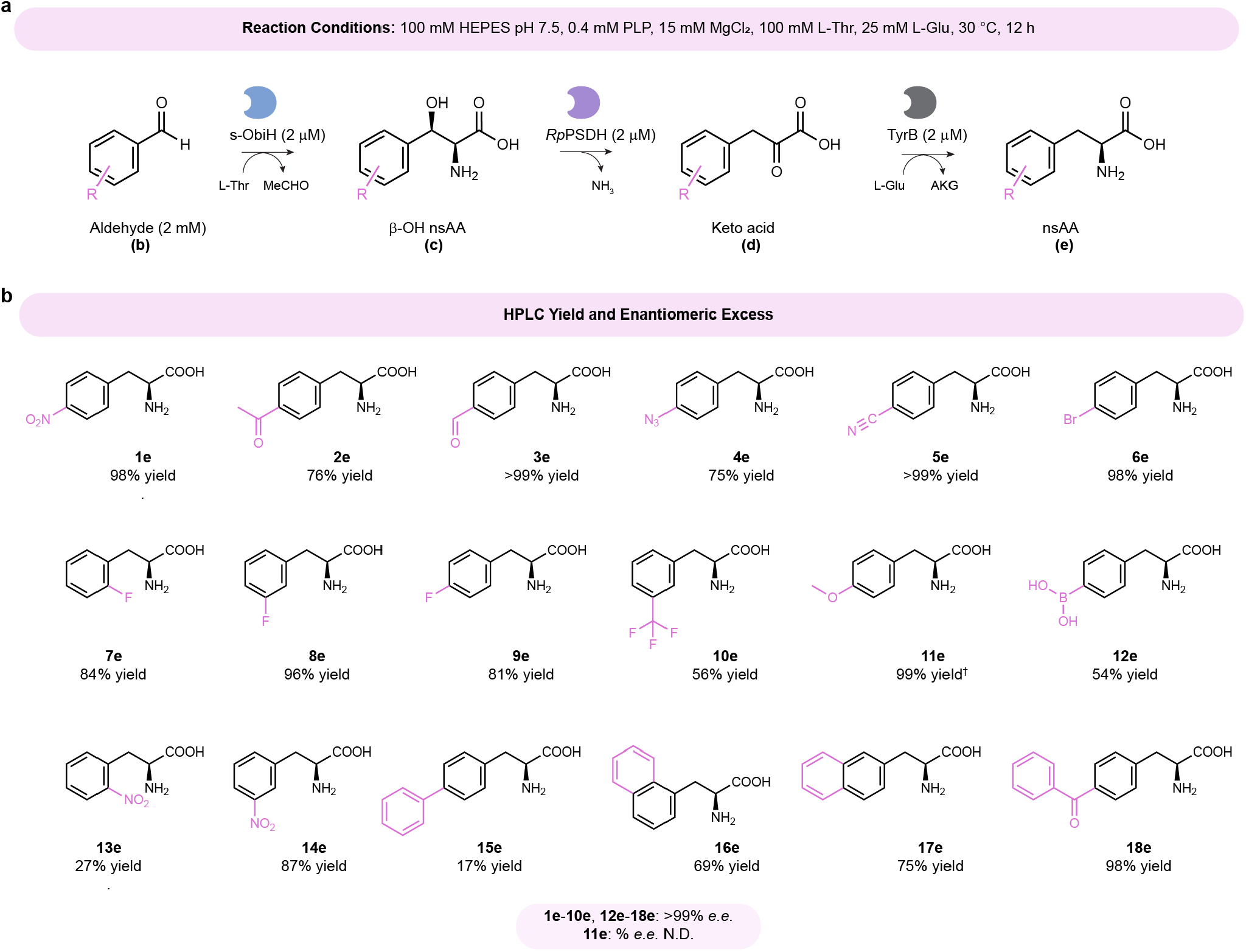
Demonstration of a broad specificity one-pot biocatalytic cascade to prepare L-phenylalanine derivatives from aryl aldehydes. (A) Reaction diagram and conditions. Three enzymes are purified and supplied at 2 μM protein concentrations: s-ObiH (0.13 mg/mL), *Rp*PSDH (0.07 mg/mL), and TyrB (0.09 mg/mL). Aldehydes are supplied at 2 mM (2% (v/v) DMSO). Other components in the reaction buffer are indicated in the top horizontal pink bar, unless otherwise specified. (B) The chemical structures and average HPLC yields of triplicate reactions are shown for 18 biosynthesized phenylalanine derivatives. We observed > 99% *e*.*e*. for all chemistries analyzed through Marfey’s analysis. The enantiomeric excess for product **11e** was not determined (N.D.). ^†^Reaction performed with 1 mM aldehyde, 5 μM s-ObiH, 2 μM *Rp*PSDH, 2 μM TyrB, pH 7.5 with 0.4 mM PLP, 15 mM MgCl, 100 mM L-Thr, 10 mM L-Glu, and 1% (v/v) DMSO at 30 °C with an endpoint of 24 h.

These reactions generated several products of interest and shed light into the preferences of the cascade. We were able to produce nsAAs used to enhance immune response^[15]^ (**1e**), as bio-orthogonal handles^[17,49]^ (**2e-4e**), in an optical probe^[50]^ (**5e**), for light-responsive biomaterial^[7]^ or protein cleavage^[51]^ (**13e**), for tuning of hydrophobic packing interactions^[52]^ (**15e-17e**), for intrinsic biological containment of organisms^[19,20]^ (**4e, 11e, 15e**), to improve protein stability^[9]^ (**18e**), for designer enzymes^[13,53]^ (**12e, 18e**), and more. Reactions with electron-withdrawing ring substituents at the *para* or *meta* positions generally performed best, with a notable exception of a reaction with electron-donating 4-methoxy group at 99% yield. The lower yields observed for **10e, 12e, 13e**, and **15e** are likely due to unfavorable electrostatics or steric hindrance for one or more pathway enzymes.

### One-pot nsAA production from carboxylic acid precursors

In pursuit of enhancing the versatility and, for some chemistries, the economic viability of our pathway, we investigated initiating our cascade from carboxylic acid precursors. Recognizing their widespread availability, reduced toxicity, lower volatility, and often lower cost compared to aldehydes, carboxylic acids presented a promising alternative. The synthesis of 4-azido-phenylalanine from its corresponding aldehyde or carboxylic acid is illustrative. The carboxylic acid is ∼33-fold less expensive per mol than the nsAA and is one of the few chemistries in which the aldehyde is more expensive compared to the nsAA (Supporting Information, Figure S22). This economic rationale underscores the potential benefits of extending the pathway to commence from a carboxylic acid.

Carboxylic acid reductases (CARs), which catalyze the conversion of acids to aldehydes, have been shown to exhibit high activity and broad substrate specificity, meeting two main criteria for our reaction cascade. Therefore, we introduced the CAR from *Segniliparus rotundus* (*Sr*CAR) into our cascade by expressing and purifying a His_6_-tagged variant. We combined 2 μM purified enzyme (0.26 mg/mL) with equimolar ratios of s-ObiH, *Rp*PSDH, and TyrB in one pot and evaluated the ability of this extended pathway to catalyze the conversion of supplemented carboxylic acids into their corresponding nsAAs over a 24-hour period (Figure 2a). The outcomes of our *in vitro* assay were highly promising, with over 75% yield observed for 10 out of the 18 different carboxylic acid precursors screened, highlighting the promiscuity of the *Sr*CAR. Additionally, four of these chemistries achieved yields exceeding 95% (Figure 2b). Notably, we were able to produce 4-azido-phenylalanine (**4e**) at 83% yield and 4-benzoyl-phenylalanine (**18e**) at 82% yield, the two chemistries we screened in our pathway in which it would be economically necessary to use a carboxylic acid precursor. However, in one case the specificity of the CAR limited reaction yields. The observed lower yield for **13a** stemmed primarily from the reduced activity of CAR on substrates featuring *ortho*-ring substituents (Supporting Information, Figure S23). Utilizing a mutant CAR engineered for improved tolerance of *ortho*-ring substituents may improve yields.^[54]^ We also confirmed production of **9e** from the **9a** precursor via MS (Supporting Information, Figure S24) but were unable to quantify yield due to overlapping peaks on the HPLC chromatogram. For our proof-of-concept reaction, we used a large excess of ATP and NADPH to ensure that the cofactor pool was not limiting. Use of co-factor regenerating enzymes or a resting whole cell catalyst platform could improve the economic viability of this cascade.^[55,56]^

**Figure 2.**
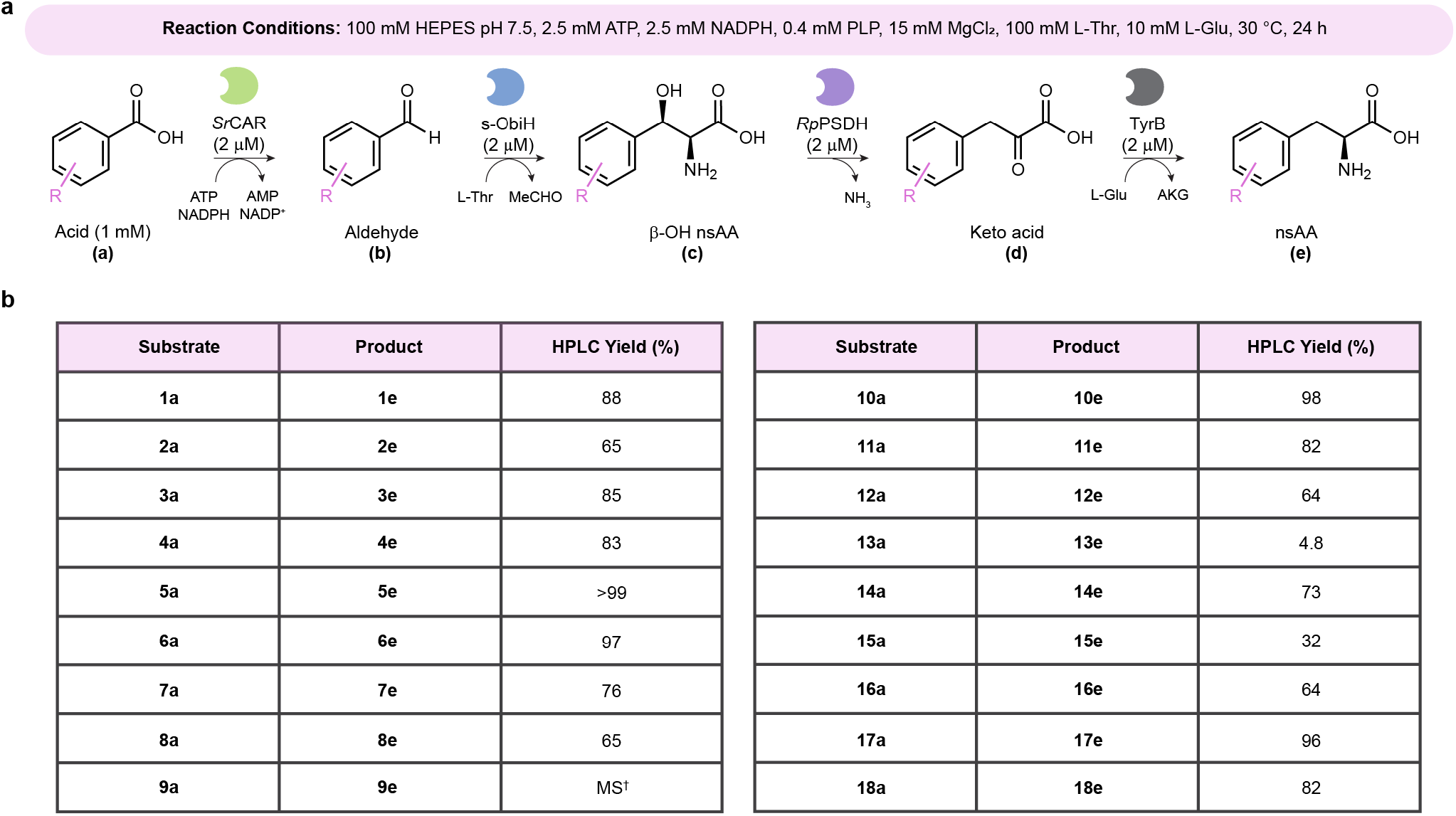
Extension of the one-pot biocatalytic cascade to include carboxylic acids as the precursor. (A) Reaction diagram that includes the carboxylic acid reductase *Sr*CAR (0.26 mg/mL) in addition to the enzymes previously supplied, all at 2 μM enzyme concentration. Carboxylic acid substrates are provided at 1 mM (1% (v/v) DMSO). Full reaction conditions are in the top horizontal. (B) Table of HPLC yields of target nsAA obtained when supplying one of 18 distinct carboxylic acids to the reaction cascade after 24 h. Reactions were performed in triplicate. ^†^Production of **9e** confirmed via MS without quantification.

### Preparative-scale synthesis of an nsAA using clarified lysate

To demonstrate the practical applicability of our cascade, we investigated the ability to harness a clarified lysate system, thus circumventing the need for costly and time-intensive downstream enzyme purification. Concurrently, we aimed to assess the activity resulting from the basal expression level of TyrB or other *E. coli* aminotransferases, potentially reducing the number of enzymes that require heterologous expression. To improve soluble expression, we utilized SUMO-tagged variants of ObiH (s-ObiH), as previously mentioned, as well as *Rp*PSDH (s-*Rp*PSDH), which were individually overexpressed in the recently engineered *E. coli* RARE.Δ16 strain.^[24]^ We chose RARE.Δ16 as our expression host, owing to the gene inactivation of 16 aldo-keto reductases and alcohol dehydrogenases, shown to effectively prevent reduction of terephthalaldehyde, which the original RARE strain could not stabilize.^[57]^ In our newly designed biocatalytic cascade, terephthalaldehyde (**3b**) can efficiently be converted to 4-formyl-L-phenylalanine (**3e**); thus, we sought to use a host for which the lysate might allow for the production of this nsAA. Encouragingly, we observed robust expression of s-ObiH and s-*Rp*PSDH (Supporting Information, Figure S25).

Initial testing of our clarified lysate-based system aimed at producing the industrially relevant nsAA for bio-orthogonal conjugation, 4-acetyl-L-phenylalanine (**2e**), from aldehyde precursors (**2b**) using a 1:1 ratio (4.5 mg/mL each) of s-ObiH and s-*Rp*PSDH clarified lysates (Figure 3a). It is important to note that *E. coli* does not natively contain ω-aminotransferases that could recognize the *para*-substituted carbonyl group of **2b** or **3b** as an amine acceptor; therefore, we did not expect those carbonyl handles for bio-orthogonal conjugation to be converted to amines by native aminotransferases present in our bacterial lysate. We observed >99% yield from 1 mM, 92% yield from 5 mM, 68% yield from 10 mM, and 46% yield from 25 mM aldehyde precursor (**2b**), as shown in Figure 3b. The reaction mixture contained 100 mM L-Thr to drive the L-TTA reaction and 25 mM L-Glu to drive the native *E. coli* aminotransferase activity inherent in the clarified lysates. Furthermore, we explored the potential for yield enhancement through co-substrate tuning, resulting in a modest 1.8-fold increase in **1e** production from 25 mM **2b** using 200 mM L-Thr and 100 mM L-Glu compared to 100 mM L-Thr and 25 mM L-Glu in a reaction containing 3 mg/mL of s-ObiH and s-*Rp*PSDH wet lysates. The increase in L-Glu concentration was the primary factor in titer increase under the conditions tested (Figure 3c). Additional targets to reduce the well-studied reversibility of the AT reaction may include the co-expression of an ornithine δ-aminotransferase while maintaining the use of L-Glu as the donor^[58]^, utilizing L-Asp as the amine donor with acetolactate synthase^[59]^, or utilizing a “smart” amine donor such as lysine^[60]^.

**Figure 3.**
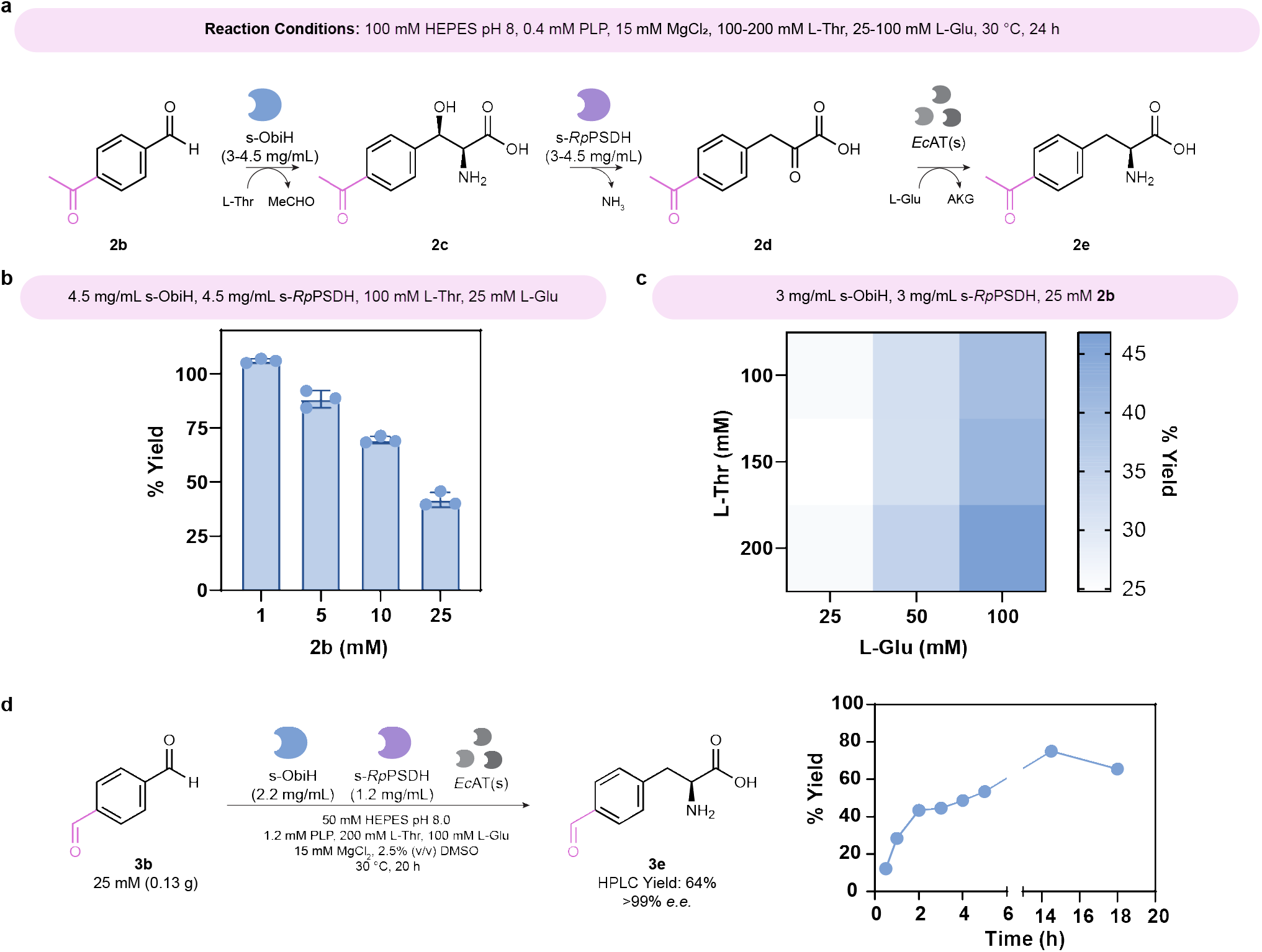
Preparative-scale nsAA synthesis using clarified lysate from the engineered *E. coli* strain RARE.Δ16. (A) Model reaction chemistry of 4-acetyl-benzaldehyde (**2b**) supplied to lysates where s-ObiH and s-*Rp*PSDH were overexpressed separately. The *E. coli* lysates contain *Ec*AT(s) at native expression levels as it is endogenous and simplifies the scale-up procedure. Reaction conditions are shown in the top horizontal bar. (B) The observed product yield of **2e** produced at different substrate loading of **2b**. Reaction performed with 1-25 mM aldehyde **2b** (0.1-2.5% (v/v) DMSO), 4.5 mg/mL s-ObiH lysate, 4.5 mg/mL *Rp*PSDH lysate, 100 mM HEPES pH 8 with 0.4 mM PLP, 15 mM MgCl_2_, 100 mM L-Thr, 25 mM L-Glu at 30 °C with an endpoint of 24 h. (C) Optimization of L-Glu amine donor for the *Ec*AT catalyzed reaction and L-Thr co-substrate for the s-ObiH-catalyzed reaction. Reaction performed with 25 mM aldehyde **2b**, 3 mg/mL s-ObiH lysate, 3 mg/mL *Rp*PSDH lysate, 100 mM HEPES pH 8 with 0.4 mM PLP, 15 mM MgCl_2_, 100-200 mM L-Thr, 25-100 mM L-Glu at 30 °C with an endpoint of 24 h. Reactions were performed in triplicate. (D) Reaction chemistry performed at 40 mL scale to convert 25 mM terephthalaldehyde (**3b**) to 4-formyl-L-phenylalanine (**3e**) under conditions shown below the reaction arrow. Final HPLC yield at 18 h was 64% at greater than 99% *e*.*e*. and time-course data is shown.

Finally, we sought to scale-up our reaction to 1 mmol scale to produce 4-formyl-L-phenylalanine (**3e**), an nsAA that could be valuable for bio-orthogonal conjugation and that has limited commercial availability. In particular, the aldehyde handle of **3e** is known to accelerate carbonyl-specific reaction times by orders of magnitude compared to the ketone handle of **2e**^[49]^, though in bacterial hosts that are not engineered to stabilize aldehydes **3e** might be prone to reduction by endogenous enzymes. By supplementing 25 mM of terephthalaldehyde (**3b**) to a 40 mL reaction containing s-ObiH and s-*Rp*PSDH clarified lysates, we observed a progressive color change from light yellow to a deeper orange as the reaction proceeded over 18 h (Supporting Information, Figure S26). The reaction reached 43% yield in just 2 h, with a final yield of 64% and greater than 99% *e*.*e*. after 18 h as determined by HPLC (Figure 3d), demonstrating an early glimpse into the scalability and efficacy of our biocatalytic approach in producing valuable nsAAs. We anticipate the system could be further improved in subsequent work by tuning concentrations of reaction components, by adding enzymes for amine donor and co-factor recycling or for acetaldehyde depletion, or by substituting cascade enzymes for homologs or engineered variants with altered specificity and activity. The one-pot biocatalytic cascade that we have devised here for the synthesis of diverse phenylalanine derivatives offers several compelling advantages over existing synthetic and biocatalytic alternatives, including other recent groundbreaking innovations in nsAA biocatalysis. For example, an unique biocatalyst that harnesses simple aryl or heterocyclic precursors for conversion to tryptophan derivatives without β-hydroxylation is the PLP-dependent enzyme TrpB.^[61]^ While an extensive evolutionary campaign sought to extend the product range of TrpB to tyrosine derivatives and succeeded in accessing two tyrosine analogs, the phenol was integral to substrate binding. This indicates barriers to the pursuit of phenylalanine derivatives that lack a *para*-hydroxy substituent without substantial modification of the active site.^[62]^ Meanwhile, recent advances at the intersection of biocatalysis and photocatalysis have leveraged the formation of quinonoid intermediates by PLP-dependent enzymes, including TrpB, L-TAs, and L-TTAs, instead for radical functionalization of the α carbon of amino acids. While these photobiocatalytic approaches have accessed several phenylalanine derivatives^[63–65]^, thus far they have relied on high catalyst concentrations as well as rare earth elements such as ruthenium or iridium or expensive organic photocatalysts. Instead, our method features low catalyst concentrations and no rare earth metals. Additionally, unlike most photobiocatalytic approaches, our system has potential for efficient function in live cells as a biochemical pathway that could be coupled directly to genetic code expansion technologies, which we explored concurrently as we were conducting this study.

The ability of these naturally occurring enzymes to catalyze apparently non-natural and sequential transformations raises the question of whether nature evolved similar biochemical pathways as alternative routes for the biosynthesis of phenylalanine derivatives that belong to natural products. To our knowledge, genes that encode characterized homologs of the enzymes featured here do not co-occur in known biosynthetic gene clusters. In *Pseudomonas fluorescens*, ObiH is part of a biosynthetic gene cluster for the synthesis of obafluorin, an antibiotic whose β-lactone core pharmacophore requires the β-hydroxy group for cyclization by the ObiD and ObiF NRPS assembly line.^[66–68]^ To look for possible homologs of *Rp*PSDH within *Pseudomonas fluorescens*, we performed a protein BLAST search using the sequence of *Rp*PSDH within the *Pseudomonas fluorescens* group taxonomy (taxid: 136843). Sequences that most closely align bear 38% identity and are annotated as *threo*-3-hydroxy-L-aspartate ammonia-lyases by the NCBI Hidden Markov Model. In general, no other phenylserine dehydratases besides *Rp*PSDH have a UniProtKB reviewed entry, and no other homologs have been reported in the literature with the exception of one PSDH from *Paraburkholderia xenovorans* LB400 DSM 17367 (*Px*PSDH), demonstrated to be active on *ortho*- and *para*-halogenated phenylserine derivatives^[69]^ and which was further used to develop a colorimetric screen for 4-(methylsulfonyl)phenylserine biosynthesis.^[70]^ Thus, it may be that other predicted phenylserine dehydratases and even predicted *threo*-3-hydroxy-L-aspartate ammonia-lyases will exhibit broad substrate scope, and it may be worth investigating whether genes encoding orthologs of these enzymes could be found in the same genomes as genes encoding L-TTA orthologs.

Finally, given the ability of *Rp*PSDH to accept β-hydroxylated leucine derivatives^[25]^, and given the ability of ObiH to accept aliphatic aldehydes as substrates^[25]^, it is highly likely that the breadth of nsAAs that can be accessed by the core of our cascade extends well beyond phenylalanine derivatives. In those cases, it may be beneficial to substitute the aminotransferase according to the specificity desired. It is also straightforward to envision the synthesis of D-amino acids instead of L-amino acids by simple replacement of the aminotransferase as the α-keto acid product of *Rp*PSDH is achiral.

## Conclusion

In summary, we have developed a one-pot biocatalytic cascade capable of synthesizing a wide range of high-value nsAAs from inexpensive and commercially available precursors. This work highlights the remarkable polyspecificity of up to four enzymes functioning together in a single pot, with surprisingly limited crosstalk between cofactors, co-substrates, or intermediates under conditions tested, resulting in high yield nsAA production. Our cascade presents a versatile method to generate industrially sought after phenylalanine derivatives, with valuable intermediates such as aldehydes (when starting from carboxylic acids; with uses in the fragrance industry), β-hydroxy nsAAs (uses in pharmaceuticals), and aromatic α-keto acids (which may serve as key precursors in drug synthesis). Additionally, we have demonstrated initial scalability of this biosynthetic pathway using clarified lysate, which allows us to leverage endogenous *E. coli* aminotransferases for catalyzing the final step in the production of an nsAA that contains a handle for bio-orthogonal conjugation. Overall, this work offers a versatile and scalable approach to reducing nsAA manufacturing costs and broadening nsAA synthesis, including for building blocks that are not commercially available.

## Acknowledgements

We sincerely thank J. Sampson in the High-Throughput Experimentation Lab at the University of Delaware for the mass spectrometry analysis and assistance that is supported by the National Institutes of Health under award P20GM104316, the National Science Foundation under award CHE-2404894, and Unidel-18D. We acknowledge support from the following funding sources: the Office of Naval Research Award No. N000142212536 (to A.M.K.); Department of Energy Joint Genome Institute under proposal 506446, a DOE Office of Science User Facility, supported by the Office of Science of the U.S. Department of Energy operated under contract no. DE-AC02-05CH11231 (to A. M. K); the National Institutes of Health Centers of Biomedical Research Excellence under grant 5P20GM104316-08 (to A. M. K.); the National Science Foundation Division of Chemical, Bioengineering, Environmental, and Transport Systems Award No. CBET-2032243 (to A.M.K. and A.P.S); and the National Institute of General Medical Sciences of the National Institutes of Health under a Chemistry-Biology Interface Training Grant T32GM133395 (to S.R.A. and M.A.J.). Additionally, we would like to thank the members of the Kunjapur group for thoughtful discussions towards this manuscript.

