## Supporting Information for "A One-Pot Biocatalytic Cascade to Access Diverse L-Phenylalanine Derivatives from Aldehydes or Carboxylic Acids"

### I. Materials and Methods

#### Strains and Plasmids

*Escherichia coli* strains and plasmids used are listed in **Table S1**. Molecular cloning and vector propagation were performed in *E. coli* DH5α. Polymerase chain reaction (PCR) based DNA replication was performed using KOD XTREME Hot Start Polymerase or the KAPA2G FAST Multiplex Kit. Cloning was performed using Gibson Assembly. Oligos for PCR amplification are shown in **Table S2**. Oligos were purchased from Integrated DNA Technologies (IDT). Expression of enzymes for purification was performed in *E. coli* BL21 (DE3). Expression of enzymes for lysate biotransformations was performed in *E. coli* MG1655 RARE.Δ16.^[1]^

#### Materials and Chemicals

The following compounds were purchased from MilliporeSigma: kanamycin sulfate (Kan), chloramphenicol (Cm), streptomycin sulfate salt (Str), dimethyl sulfoxide (DMSO), potassium phosphate dibasic, potassium phosphate monobasic, magnesium sulfate, calcium chloride dihydrate, glycerol, Tris base, glycine, HEPES, ATP, and KOD XTREME Hot Start polymerase. The following were purchased from TCI America: D-glucose, phenylpyruvic acid. 4-nitrobenzoic acid (1a), 4-acetylbenzoic acid (2a), 4-azidobenzoic acid (4a), 3-(trifluoromethyl)benzoic acid (10a), 2-nitrobenzoic acid (13a), 3-nitrobenzoic acid (14a), 2-naphthoic acid (17a), terephthalaldehyde (3e), 2-nitrobenzaldehyde (13b), 3-nitrobenzaldehyde (14b), biphenyl-4-carboxaldehyde (15b), 1-naphthaldehyde (16b), 2-fluoro-L-phenylalanine (7e), 3-fluoro-L-phenylalanine (8e), and O-methyl-L-tyrosine (16e). The following were purchased from Peptech: 4-nitro-L-phenylalanine (1e), 4-acetyl-L-phenylalanine (2e), 4-cyano-L-phenylalanine (5e), 4-bromo-L-phenylalanine (6e), 4-fluoro-L-phenylalanine (9e), 2-nitro-L-phenylalanine (13e), 3-(1-Naphthyl)-L-alanine (16e), 3-(2-Naphthyl)-L-alanine (17e), and 4-benzoyl-L-phenylalanine (18e). The following were purchased from Thermo Scientific Chemicals: FDAA (Marfey's Reagent) (1-fluoro-2-4-dinitrophenyl-5-L-alanine amide, benzoic acid, 4-bromobenzoic acid (6a), 2-fluorobenzoic acid (7a), 3-fluorobenzoic acid (8a), 4-fluorobenzoic acid (9a), 1-naphthoic acid (16a), 4-benzoylbenzoic acid (18a), 4-bromobenzaldehyde (6b), 3-fluorobenzaldehyde (8b), 4-fluorobenzaldehyde (9b), and 3-nitro-L-phenylalanine (14e). The following were purchased from Sigma-Aldrich: benzaldehyde, terephthalic acid (3a), 4-methoxybenzoic acid (11a), biphenyl-4-carboxylic acid (15a), 4-nitrobenzaldehyde (1b), 4-formylbenzonitrile (5b), 4-anisaldehyde (11a), 4-formylphenylboronoic acid (12b), 2-naphthaldehyde (17b), and L-phenylalanine. Agarose, ethanol, 2-fluorobenzaldehyde (7b), L-glutamic acid monopotassium salt monohydrate, and L-threonine were purchased from Alfa Aesar. Acetonitrile, trifluoroacetic acid (TFA), sodium chloride, LB Broth powder (Lennox), and LB Agar powder (Lennox) were purchased from Fisher Chemical. Taq DNA ligase was purchased from GoldBio. L-glutamic acid, 3-(trifluoromethyl)benzaldehyde (10b), and 4-borono-L-phenylalanine (12e) were purchased from ACROS Organics. (S)-2-Amino-3-(3-(trifluoromethyl)phenyl)propanoic acid (10e), 4-boronobenzoic acid (12a), and (S)-2-Amino-3-(3-(trifluoromethyl)phenyl)propanoic acid (10e) were purchased from Ambeed. 4-cyanobenzoic acid (5a), 4-azidobenzaldehyde (4b), and 4-carboxy-L-phenylalanine (19e) were purchased from ChemCruz. 4-azido-L-phenylalanine (4e) was purchased from Bachem. Biphenylalanine (15e) was purchased from Combi-blocks. 4-nitrophenypyruvic acid was purchased from abcr GmbH. (2S)-2-amino-3-(4-formylphenyl)propanoic acid hydrochloride (3e) was custom chemically synthesized by ChiroBlock GmbH. Phusion DNA polymerase and T5 exonuclease were purchased from New England BioLabs (NEB). Sybr Safe DNA gel stain was purchased from Invitrogen. NADPH (tetrasodium salt) and 4-acetylbenzaldehyde (2b) were purchased from Santa Cruz Biotechnology. Anhydrotetracycline (aTc) was purchased from Cayman Chemical. KAPA2G FAST Multiplex Kit was purchased from Roche.

#### Culture Conditions

Cultures were grown in LB-Lennox medium (LBL: 10 g/L bacto tryptone, 5 g/L sodium chloride, 5 g/L yeast extract) or 2xYT medium (16 g/L tryptone, 10 g/L yeast extract, 5 g/L sodium chloride). Cultures were inoculated from an overnight culture and grown in appropriate antibiotic (30 µg/mL Kan) at 37 °C with shaking at 250 RPM in a 1 L baffled shake flask. At an OD_600_= 0.5-0.8, 0.2 µM anhydrotetracycline (aTc) added to induce enzyme expression. Cultures were incubated for an additional 5 h at 30 °C, followed by incubation at 18 °C for 16 h with shaking at 250 RPM unless otherwise specified.

#### HPLC and LC-MS analysis

Metabolites of interest were quantified via high-performance liquid chromatography (HPLC) using an Agilent 1100 Infinity model equipped with a Zorbax Eclipse Plus-C18 column (part number: 959701-902, 5 µm, 95Å, 2.1 x 150 mm). To quantify compounds of interest, an initial mobile phase of solvent A/B = 95/5 was used (solvent A, water, 0.1% TFA; solvent B, acetonitrile, 0.1% TFA) and maintained for 5 min. A gradient elution was performed (A/B) with: gradient from 95/5 to 50/50 for 5-12 min, gradient from 50/50 to 0/100 for 12-13 min, gradient from 0/100 to 95/5 for 13-14, and equilibration at 95/5 for 14-15 min. A flow rate of 1 mL min^-1^ was maintained, and absorbance was monitored at 210, 250, 270, 280 and 300 nm. Confirmation of product production was performed using a Waters AQUITY Arc UPLC H-Class with a diode array coupled to a Waters AQUITY QDa Mass Detector. Product was analyzed using a Waters Cortecs UPLC C18 column with an initial mobile phase of solvent A/B = 95/5 (solvent A, water, 0.1% formic acid; solvent B, acetonitrile, 0.1% formic acid) and maintained for 5 min. A gradient elution was performed (A/B) with: gradient from 95/5 to 10/90 for 5-7 min, an isocratic flow at 10/90 for 7-10 min, then gradient from 10/90 to 95/5 for 10-10.5 min and a final isocratic step at 95/5 for 10-12 min. A flow rate of 1 mL min^-1^ was maintained.

#### Cloning, Expression and Purification of Pathway Enzymes

Molecular cloning and vector propagation were performed in *E. coli* DH5α (NEB). PSDH gene from *Ralstonia pickettii* (*Rp*PSDH) was purchased as a gene fragment from Twist Bioscience and codon optimized for expression in *E. coli*. All primers were ordered from IDT. The AT gene from *Escherichia Coli* (TyrB) was PCR amplified from the *E. coli* genome using KAPA2G HotStart DNA polymerase. For the L-TTA, the L-TTA gene from *Pseudomonas fluorescens* (ObiH) on a pZE vector harboring an N-terminal-SUMO-His_6_-tag for purification and improved solubility (s-ObiH), kanamycin resistance gene for antibiotic selection, and ColE1 ori from Jones, *et. al* was used.^[2]^ For the PSDH and AT, amplified fragments were then cloned into a PCR-amplified pZE vector harboring an N-terminal-His_6_-tag for purification, kanamycin resistance gene, and ColE1 ori. The PSDH was also cloned into a PCR-amplified pZE vector harboring an N-terminal-SUMO-His_6_-tag for purification and improved solubility, kanamycin resistance gene, and ColE1 ori. For the CAR, the CAR gene from *Segniliparus rotundus* (*Sr*CAR) on a pZE vector harboring an N-terminal His_6_-tag for purification, kanamycin resistance gene, ColE1 ori, and a recombinant 4’-phosphopantetheinyl transferase gene (*sfp*) from *Bacillus subtilis* (*Bs*Sfp) from Gopal, *et. al* was used.^[3]^ Plasmids were verified by Sanger sequencing.

For expression, each plasmid encoding expression of each pathway enzyme was transformed into *E. coli* BL21(DE3) (NEB). 300 mL of LB in 1 L baffled flasks were supplemented with 30 µg/mL Kan and inoculated from an overnight culture at a 1:100 inoculation ratio and incubated at 37 °C with shaking at 250 RPM. Cultures were induced at mid-exponential phase (OD_600_=0.5-0.8) with 0.2 µM aTc, and then incubated 30 °C for 5 h, followed by overnight expression at 18 °C. Cells were harvested by centrifugation.

For purification of s-ObiH and *Rp*PSDH, cells were resuspended in Lysis buffer (25 mM HEPES pH 7.4, 250 mM NaCl, 0.4 mM PLP, 10 mM MgCl_2_ and 10 mM imidazole). The cells were lysed by sonication followed by centrifugation at 18,213 × *g* for 1 h. The supernatant was sterile filtered through a 0.22 µm syringe filter and purified using an Ni-Sepharose affinity chromatography (HisTrap HP, 5 mL) via an AKTA Pure fast protein liquid chromatography (FPLC) system using Lysis Buffer A (25 mM HEPES pH 7.4, 250 mM NaCl, 0.4 mM PLP, 10 mM MgCl_2_, and 10 mM imidazole) and Elution Buffer B (25 mM HEPES pH 7.4, 250 mM NaCl, 0.4 mM PLP, 10 mM MgCl_2_, and 250 mM imidazole). The column was equilibrated with 2.5% Buffer B and after performing an isocratic 12% Buffer B (40 mM) and 18% Buffer B (54 mM imidazole) wash, elution was performed at 250 mM imidazole. Purified fractions were pooled, concentrated, and dialyzed against dialysis buffer (100 mM HEPES pH 7.4, 300 mM NaCl, 0.4 mM PLP, and 10 mM MgCl_2_) with a 30 kDa molecular weight cutoff centrifugal filter (Amicon Ultra, Millipore). Samples were then flash frozen in Eppendorf tubes in an ultracold ethanol bath and stored at -80 ºC.

For TyrB and *Sr*CAR purification, cells were resuspended in Lysis buffer (25 mM HEPES pH 7.4, 250 mM NaCl, and 10 mM imidazole). The cells were lysed by sonication followed by centrifugation at 18,213 × *g* for 1 h. The cleared supernatant was sterile filtered through a 0.22 μm syringe filter and purified using an Ni-Sepharose affinity chromatography (HisTrap HP, 5 mL) via an AKTA Pure fast protein liquid chromatography (FPLC) system using Lysis Buffer A (25 mM HEPES pH 7.4, 250 mM NaCl, and 10 mM imidazole) and Elution Buffer B (25 mM HEPES pH 7.4, 250 mM NaCl, and 250 mM imidazole). The column was equilibrated with 2.5% Buffer B and after performing an isocratic 12% Buffer B (40 mM) and 18% Buffer B (54 mM imidazole) wash, elution was performed at 250 mM imidazole. Purified fractions were pooled, concentrated, and dialyzed against dialysis buffer (100 mM HEPES pH 7.4, 300 mM NaCl, and 10 mM MgCl_2_) with a 30 kDa molecular weight cutoff centrifugal filter (Amicon Ultra, Millipore). Samples were then flash frozen in Eppendorf tubes in an ultracold ethanol bath and stored at -80 ºC.

#### *In vitro* PSDH Assay

Reactions were run at 100 µL scale in triplicate in 96-well plates, unless otherwise specified. The reaction was performed with 2 µM *Rp*PSDH (0.07 mg/mL), 100 mM HEPES pH 7.5, 1 mM phenylserine (100 mM stock in water), 400 µM PLP, and 15 mM MgCl_2_ in both the absence and presence of 100 mM L-Thr for 2 h at 30 ºC with shaking in a Thermo Scientific plate shaker (catalog number 88882006) at 1000 RPM. Samples were quenched with 1% TFA. Samples were chilled at -80 ºC for at least 1 h prior to centrifugation to clear insoluble protein precipitate. Samples were then analyzed by HPLC-UV.

#### In vitro AT Assay

The reaction was performed with 2 µM TyrB (0.09 mg/mL), 100 mM HEPES pH 7.5, 400 µM PLP, 15 mM MgCl_2_, 10 mM L-Glu, and 0.5 mM phenylpyruvate (100 mM stock in 70% ethanol) or 0.5 mM 4-nitro-phenylpyruvate (100 mM stock in 70% ethanol) for 15 min at 30 ºC at 1000 RPM. Samples were quenched with 1% TFA. Samples were chilled at -80 ºC for at least 1 h prior to centrifugation to clear insoluble protein precipitate. Samples were then analyzed by HPLC-UV.

#### In vitro one-pot TTA, PSDH, AT Assay

The reaction was performed with 2 µM s-ObiH (0.13 mg/mL), 2 µM *Rp*PSDH (0.07 mg/mL), 2 µM TyrB (0.09 mg/mL), 100 mM HEPES pH 7.5, 400 µM PLP, 15 mM MgCl_2_, 100 mM L-Thr, and 25 mM L-Glu. For the one-pot production of phenylalanine, 2 mM of each substrate and possible intermediate (benzaldehyde, L-*threo*-phenylserine, or phenylpyruvic acid) was added to the reaction mixture and quenched with 1% TFA after 0.5 h at 30 ºC at 1000 RPM. For the production of diverse phenylalanine derivatives, 2 mM of each aldehyde tested (100 mM stock in DMSO) was added to the reaction mixture and subsequently incubated for 12 h at 30 ºC at 1000 RPM. Samples were quenched at a 4x dilution in 3:1 v/v ratio of methanol:1M HCl. Samples were chilled at -80 ºC for at least 1 h prior to centrifugation to clear insoluble protein precipitate. Samples were then analyzed by HPLC-UV.

#### In vitro one-pot CAR, TTA, PSDH, AT Assay

The reaction was performed with 2 µM SrCAR (0.26 mg/mL), 2 µM s-ObiH (0.13 mg/mL), 2 µM *Rp*PSDH (0.07 mg/mL), 2 µM TyrB (0.09 mg/mL), 100 mM HEPES pH 7.5, 2.5 mM ATP, 2.5 mM NADPH, 400 µM PLP, 15 mM MgCl_2_, 100 mM L-Thr, 10 mM L-Glu, 1% (v/v) DMSO, and 1 mM of acid tested (100 mM stock in DMSO) for 24 h at 30 ºC at 1000 RPM. Samples were quenched with 1% TFA. Samples were chilled at -80 ºC for at least 1 h prior to centrifugation to clear insoluble protein precipitate. Samples were then analyzed by HPLC-UV.

#### Preparation of s-ObiH and s-RpPSDH Cell-free Extracts

5 mL starter cultures of RARE.∆16 pZE-s-ObiH and RARE.∆16 pZE-s-*Rp*PSDH started from glycerol stocks were grown in LB media with 30 µg/mL Kan at 37 °C with shaking overnight until the cultures reached saturation. Then, 250 mL of 2xYT media with 30 µg/mL Kan in a 1 L baffled flask was inoculated from the overnight culture at a 1:100 inoculation ratio and incubated at 37 °C with shaking at 250 RPM. Cultures were induced at OD_600_=0.5-0.8 with 0.2 µM aTc and then incubated at 20 °C with shaking at 250 RPM for 24 h. Cells were harvested by centrifugation.

Cells were washed once in 25 mL of 25 mM HEPES pH 8 and then resuspended in a cold lysis buffer (25 mM HEPES pH 8) at 5 mL lysis buffer per g cell pellet and sonicated using a QSonica Q125 sonicator with cycles of 5 s at 90% amplitude and 10 s off for 8 min. Crude lysates were centrifuged at 17000 × *g* for 30 min.

#### Clarified Lysate Assays for Synthesis of 4-acetylphenylalanine

The total protein concentrations of the s-ObiH and s-*Rp*PSDH clarified lysates were measured by Bradford assay using bovine serum albumin as a reference. The reaction was performed with 3-4.5 mg/mL s-ObiH lysate and 3-4.5 mg/mL s-*Rp*PSDH lysate in reaction buffer containing 100 mM HEPES pH 8, 400 µM PLP, 15 mM MgCl_2_, 100-200 mM L-Thr, 25-100 mM L-Glu, 0.1-2.5% (v/v) DMSO, and 1-25 mM of 4-acetyl-benzaldehyde (**2b**) (1 M stock in DMSO) for 24 h at 30 ºC at 1000 RPM. Samples were quenched with a 3:1 v/v ratio of methanol:2M HCl. Samples were chilled at -80 ºC for at least 1 h prior to centrifugation to clear insoluble protein precipitate. Samples were then analyzed by HPLC-UV.

#### Preparative Scale Reaction for Synthesis of 4-formylphenylalanine

The soluble lysates were dialyzed using 3.5 kDa MWCO Spectra/Por 3 Dialysis Tubing against a dialysis buffer containing 50 mM HEPES pH 7.5, 0.2 mM pyridoxal-5’-phosphate monohydrate (PLP), 5 mM MgCl_2_, and 150 mM NaCl. Dialysis was performed at 4 ºC with gentle stirring for 18 h. Lysate total protein concentrations were measured after dialysis by Bradford assay using bovine serum albumin as a reference.

To a 50 mL round bottom flask was added 26.7 mL DI water, 0.477 g HEPES (final concentration: 50 mM), 0.057 g MgCl_2_ (final concentration: 15 mM), 0.012 g PLP (final concentration: 1.2 mM), 0.953 g L-threonine (final concentration: 200 mM), and 0.813 g L-glutamic acid monopotassium salt monohydrate (final concentration: 100 mM). The reaction buffer was titrated to a pH of 8. s-ObiH lysate (final lysate concentration: 2.2 mg/mL) s-*Rp*PSDH lysate (final lysate concentration 1.2 mg/mL) was then added to the reaction buffer. The reaction was gently stirred at 200 RPM using a stir bar and the temperature was maintained at 30 ºC. 0.13 g of terephthalaldehyde (final concentration: 25 mM) dissolved in 1 mL of DMSO (final concentration 2.5% v/v) was added to initiate the reaction at 40 mL scale. Timepoints were taken at 30 min, 1 h, 2 h, 3 h, 4 h, 5 h, 14.5 h, and 18 h by quenching 4 µL of the reaction in 196 µL of 3:1 v/v ratio of methanol:1M HCl (50x dilution). The samples were vigorously mixed and centrifuged at 18,213 × *g* for 1 minute. Samples were analyzed by HPLC-UV.

#### Enantiomeric Excess by Marfey’s Derivatization

To a 200 µL PCR tube was added 20 µL of 1 M sodium bicarbonate, 25 µL of 10 mM Marfey’s reagent (prepared in acetonitrile), and 5 uL of quenched reaction (previous diluted 4-fold in methanol:1M HCl). The reaction was incubated at 40 ºC for 20 h in a thermal heating block. The reactions were quenched by addition 50 µL of 4:1 (v/v) mixture of acetonitrile:2 M HCl. Degassed samples were immediately loaded onto a Waters AQUITY Arc UPLC H-Class equipped with a diode array coupled to a Waters AQUITY QDa Mass Detector. Separation was achieved using a Waters Cortecs C18 column with solvent A, water, 0.1% formic acid; solvent B, acetonitrile, 0.1% formic acid. For method 1: a gradient elution was performed (A/B) with the following method: gradient from 90/10 to 40/60 for 5 min, a gradient from 40/60 to 10/90 for 5-14 min, an isocratic flow at 10/90 for 14-14.5 min, then gradient from 10/90 to 90/10 for 14.51-15 min. For method 2: a gradient from 90/10 to 20/80 for 14 min, then gradient from 20/80 to 90/10 for 14-14.5 min. For method 3: a gradient from 90/10 to 60/40 for 14 min, then gradient from 60/40 to 90/10 for 14-14.5 min. A flow rate of 1 mL min^-1^ was maintained, and absorbance was monitored at 340 nm.

### II. Supplementary Tables

#### Table S1. Strains and Plasmids used in this study.

| Name | Relevant genotype | Source |
| --- | --- | --- |
| E. coli strains |  |  |
| DH5α | F– Φ80*lac*ZΔM15 Δ(l*ac*ZYA-argF) U169 *rec*A1 *end*A1 *hsd*R17 (rK–, mK+) *pho*A *sup*E44 λ– *thi*-1 *gyr*A96 *rel*A1 | NEB |
| RARE. Δ16 | MG1655(DE3) ∆dkgB ∆yeaE ∆(yqhC-dkgA) ∆yahK ∆yjgB ∆adhP ∆fucO ∆eutG ∆yiaY ∆adhE ∆eutE ∆gldA ∆gpr ∆ybbO ∆yghA | Previous study^[1]^ |
| BL21 (DE3) | *fhu*A2 [lon] *omp*T *gal* (λ DE3) [dcm] ∆*hsd*S | NEB |
| 1 | BL21 (DE3) harboring pZE-s-ObiH | Previous study^[2]^ |
| 2 | BL21 (DE3) harboring pZE-*Rp*PSDH | This study |
| 3 | BL21 (DE3) harboring pZE-s-*Rp*PSDH | This study |
| 4 | BL21 (DE3) harboring pZE-TyrB | This study |
| 5 | BL21 (DE3) harboring pZE-*Sr*Car-sfp | Previous study^[3]^ |
| 6 | BL21 (DE3) harboring pZE-s-*Pb*TTA | Previous study^[2]^ |
| 7 | RARE.Δ16 (DE3) harboring pZE-s-ObiH | This study |
| 8 | RARE.Δ16 (DE3) harboring pZE-s-*Rp*PSDH | This study |
| Plasmids | | |
| pZE-s-ObiH | pZE plasmid harboring a codon optimized L-threonine transaldolase from *Pseudomonas fluorescens* (ObiH) containing an N-terminal His_6_ and SUMO tag (denoted s-ObiH). ColE1 Ori, Kan^R^, TetR, Tet promoter | Previous study^[2]^ |
| pZE-*Rp*PSDH | pZE plasmid harboring a codon optimized phenylserine dehydratase from *Ralstonia pickettii* PS22 (*Rp*PSDH) containing an N-terminal His_6_ tag. ColE1 Ori, Kan^R^, TetR, Tet promoter | This study |
| pZE-s-*Rp*PSDH | pZE plasmid harboring a codon optimized phenylserine dehydratase from *Ralstonia pickettii* PS22 (*Rp*PSDH) containing an N-terminal His_6_ and SUMO tag (denoted s-*Rp*PSDH). ColE1 Ori, Kan^R^, TetR, Tet promoter | This study |
| pZE-TyrB | pZE plasmid harboring an aminotransferase from *Escherichia Coli* (TyrB) containing an N-terminal His_6_ tag. ColE1 Ori, Kan^R^, TetR, Tet promoter | This study |
| pZE-*Sr*CAR-sfp | pZE plasmid harboring a codon optimized carboxylic acid reductase from *Segniliparus rotundus* (*Sr*CAR) and a codon optimized phosphopantetheinyl transferase from *Bacillus* *subtilis* (sfp). ColE1 Ori, Kan^R^, TetR, Tet promoter | Previous study^[3]^ |
| pZE-s-*Pb*TTA | pZE plasmid harboring a codon optimized L-threonine transaldolase from *Parachlamydiales bacterium* (*Pb*TTA) containing an N-terminal His_6_ and SUMO tag (denoted s-*Pb*TTA). ColE1 Ori, Kan^R^, TetR, Tet promoter | Previous study^[2]^ |

#### Table S2. Oligonucleotides used in this study.

| Oligo Name | Sequence |
| --- | --- |
| TyrB pZE ins Rev | GGGATCCCCCATCAAGTTACATCACCGCAGCAAAC |
| TyrB pZE ins Fwd | CCATCACCATCATCACCACGTGTTTCAAAAAGTTGACGCC |
| pZE His bbone Fwd | TAACTTGATGGGGGATCCC |
| pZE His bbone Rev | GTGGTGATGATGGTGATGG |
| pZE-RpPSDH bbone Fwd | CGCGTCACCTTGGGCTGGGATAACTTGATGGGGGATCCC |
| pZE-RpPSDH bbone Rev | GTAGTGTCCAGCTGGGTCATGTGGTGATGATGGTGATGG |
| pZE-His-SUMO-RpPSDH bbone Fwd | CGCGTCACCTTGGGCTGGGATAACTTGATGGGGGATCC |
| pZE-His-SUMO-RpPSDH bbone Rev | GTAGTGTCCAGCTGGGTCATGCCCTGAAAATACAGATTTTCT |

Table S3. Twist gene fragment for cloning in this study. The start codon is underlined.

| Oligo Name | GenBank Number | Sequence |
| --- | --- | --- |
| *Rp*PSDH | BAC53614.1 | ATGACCCAGCTGGACACTACCACACTTCCGGACTTATCTGCCATTGCTGGGTTACGCGCTCGCTTGAAACAATGGGTTCGCACCACTCCAGTGTTCGACAAGACCGATTTCGAACCTGTCCCAGGAACCGCTGTGAATTTCAAATTAGAGCTTCTGCAGGCGAGCGGGACATTTAAGGCACGCGGGGCGTTTTCTAATTTGTTAGCCCTTGACGACGATCAGCGCGCAGCCGGTGTTACCTGCGTGTCCGCAGGAAATCATGCAGTTGGTGTGGCATACGCTGCGATGCGCCTTGGCATCCCGGCAAAGGTTGTGATGATTAAGACCGCATCGCCAGCGCGCGTGGCGCTTTGCCGTCAATATGGGGCTGAAGTAGTTCTGGCGGAAAACGGCCAAACCGCTTTTGACACGGTCCACCGTATCGAATCAGAGGAGGGACGCTTTTTCGTACATCCCTTTAACGGTTATCGTACTGTTCTTGGGACCGCAACATTAGGACATGAATGGTTAGAGCAGGCTGGGGCGTTGGATGCGGTGATTGTACCCATCGGTGGCGGTGGACTTATGGCGGGGGTCTCGACAGCAGTAAAGTTGTTAGCCCCGCAGTGTCAGGTGATCGGGGTGGAACCAGAGGGAGCAGATGCTATGCATCGCTCTTTCGAAACTGGGGGTCCTGTGAAGATGGGCTCGATGCAAAGTATCGCCGACAGCTTAATGGCGCCTCATACTGAGCAGTATTCGTATGAATTGTGCCGCCGTAACGTGGATCGCTTGGTAAAGGTGTCCGACGATGAGCTTCGCGCAGCAATGCGTCTGCTGTTTGACCAACTGAAACTTGCCACAGAGCCCGCATGTGCAACAGCGACTGCTGCTTTAGTCGGCGGCTTAAAAGCAGAATTGGCGGGAAAGCGCGTCGGCGTGTTACTTTGCGGTACAAATACGGATGCAGCTACGTTTGCGCGTCACCTTGGGCTGGGA |

### III. Supplementary Figures

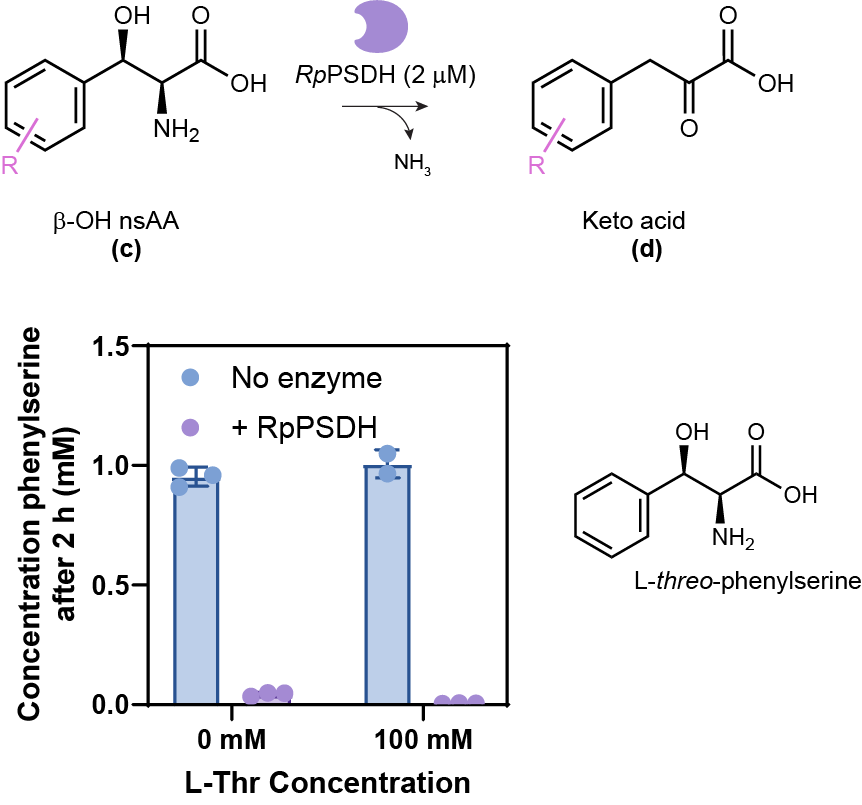

Figure S1. Evaluation of the PSDH-catalyzed pathway step. HPLC measurement of endpoint (2 h) concentrations of the keto acid product resulting from *Rp*PSDH enzyme activity on L-*threo*-phenylserine (1 mM), in the absence or presence of 100 mM L-Thr, a co-substrate of the reaction catalyzed by L-TTA.

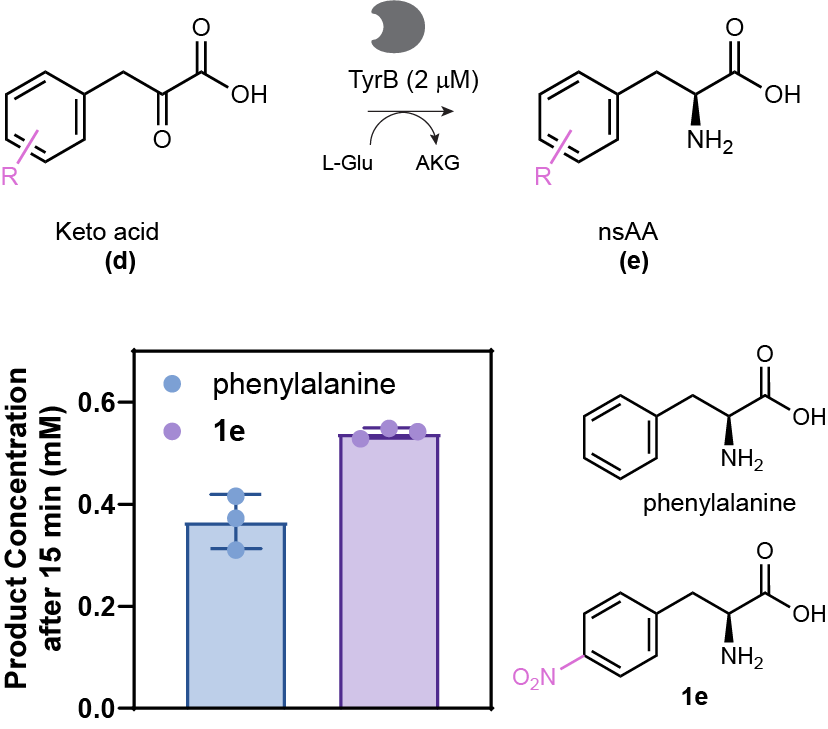

Figure S2. Evaluation of the AT-catalyzed reaction step. HPLC measurement of endpoint (15 min) concentrations of phenylalanine and **1e**. The final product demonstrates the ability of the TyrB enzyme to convert a native substrate (phenylpyruvate, 0.5 mM) and a non-native substrate (4-nitro-phenylpyruvate, 0.5 mM) to their associated α-amino acids in the presence of 10 mM L-Glu as the amine donor.

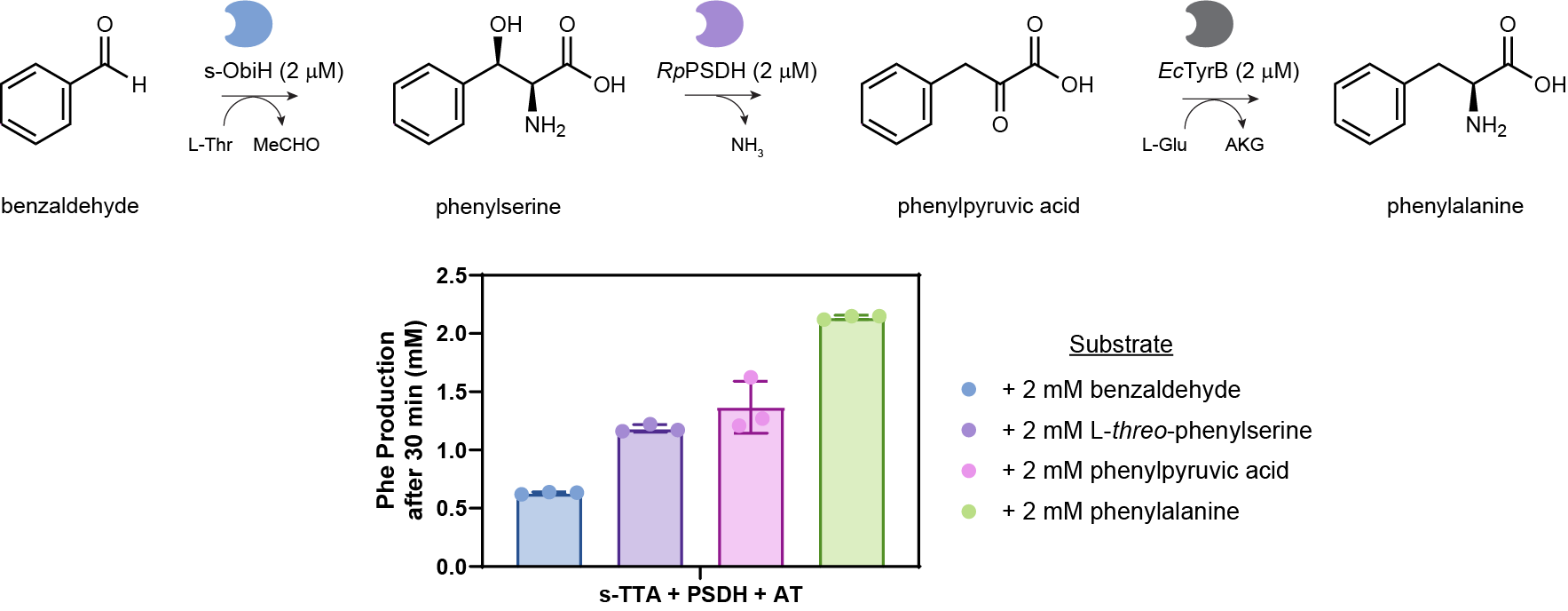

Figure S3. Compatibility of the three purified enzymes assembled in a one-pot reaction for a multi-step biocatalytic cascade. The one-pot reaction was prepared by adding equimolar enzyme concentration (2 µM) of each purified enzyme (s-ObiH (0.13 mg/mL), *Rp*PSDH (0.07 mg/mL), and TyrB (0.09 mg/mL)) to a reaction mixture containing 0.4 mM PLP, 15 mM MgCl_2_, 100 mM L-Thr, and 25 mM L-Glu. Each possible intermediate was supplied to test enzyme activity of each step in the assembled mixture. Benzaldehyde is used as a model chemistry as the L-TTA (s-ObiH) converts it to phenylserine, which then lead to the native substrates for the PSDH (*Rp*PSDH) and AT (TyrB) catalyzed reactions. The concentration of phenylalanine, the desired final product, is measured by HPLC 30 min after reaction initiation upon supplementing 2 mM of one of the respective substrates (benzaldehyde, phenylserine, or phenylpyruvate) or 2 mM of the product phenylalanine to confirm product stability.

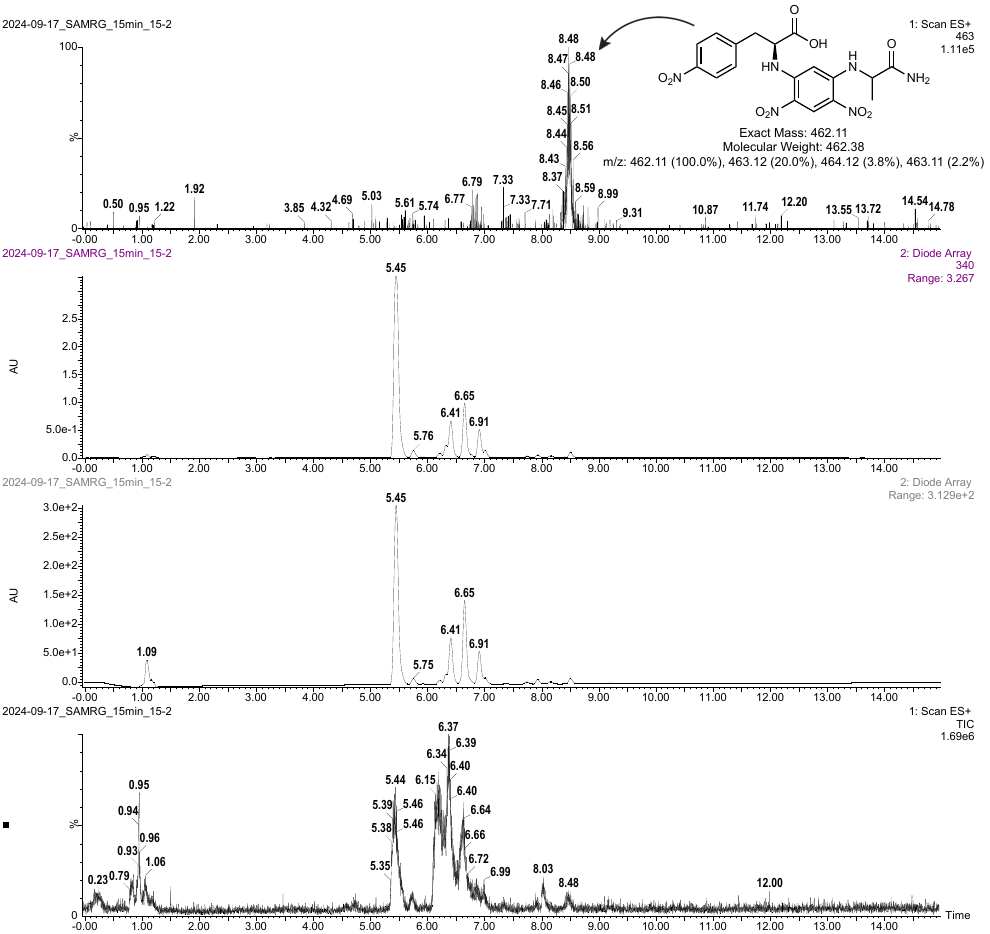

Figure S4. Marfey’s analysis on the one-pot s-ObiH, *Rp*PSDH, TyrB catalyzed reaction with **1b**. LC-MS method 2 was used for this analysis.

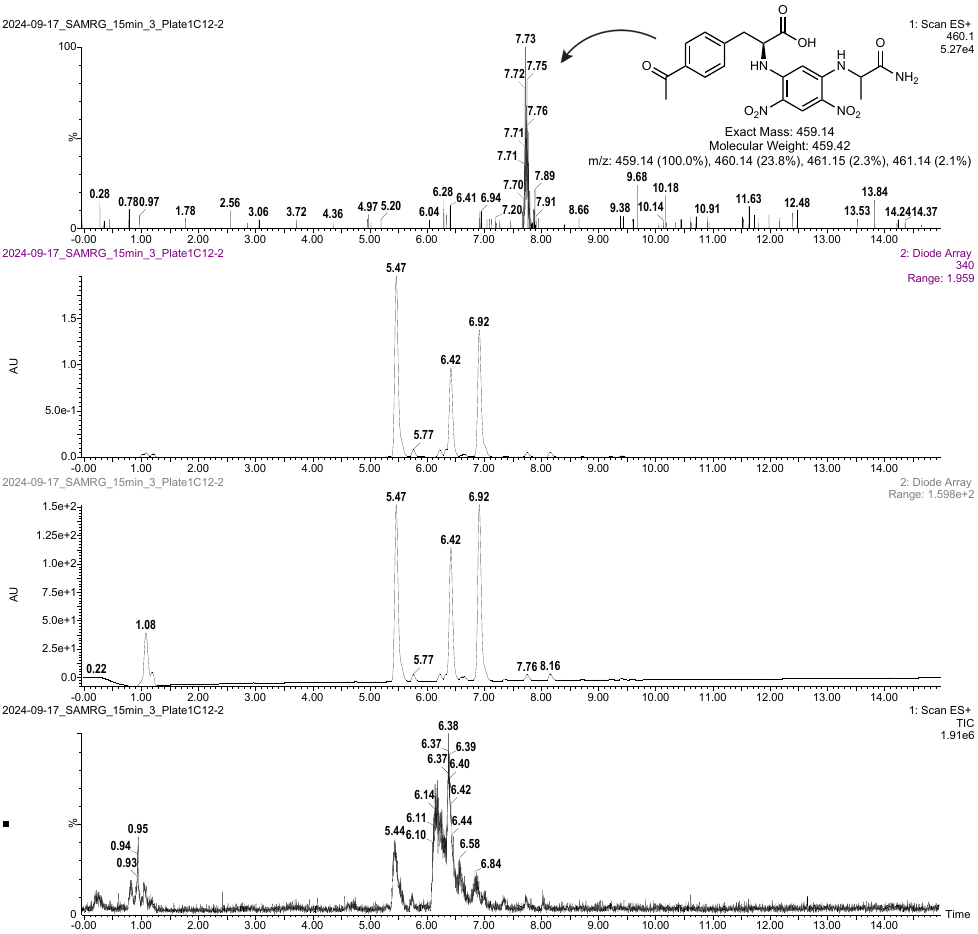

Figure S5. Marfey’s analysis on the one-pot s-ObiH, *Rp*PSDH, TyrB catalyzed reaction with **2b**. LC-MS method 2 was used for this analysis.

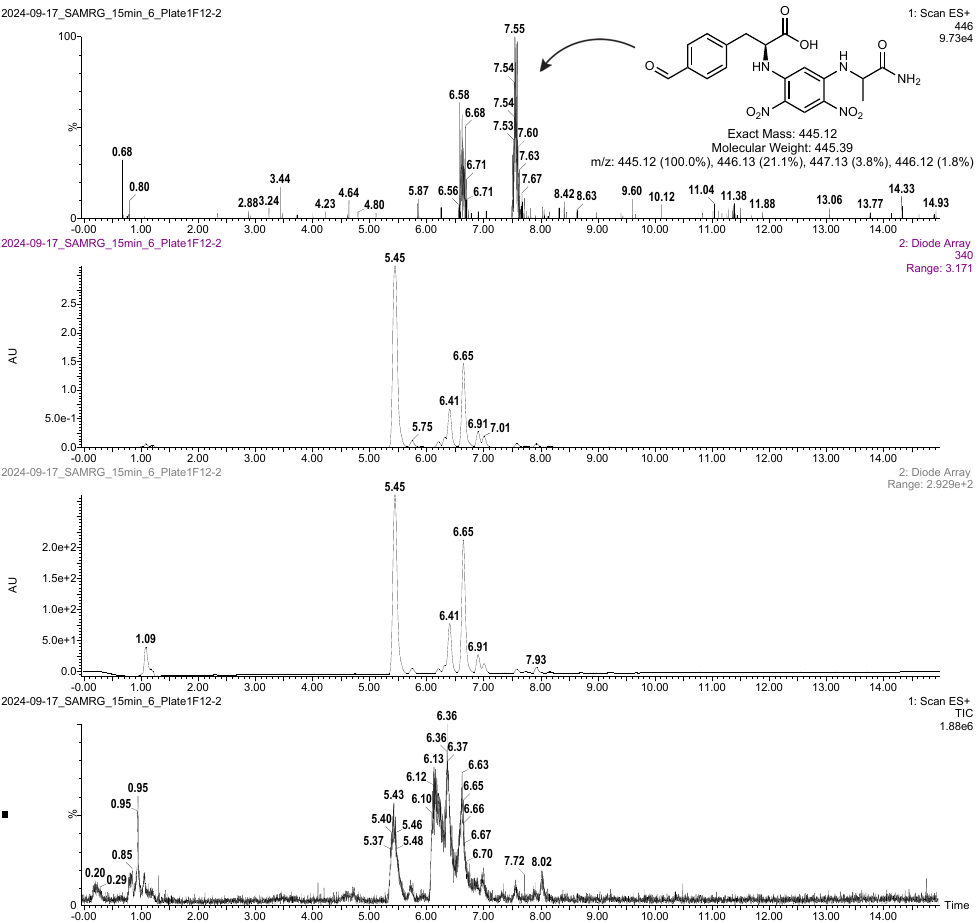

Figure S6. Marfey’s analysis on the one-pot s-ObiH, *Rp*PSDH, TyrB catalyzed reaction with **3b**. The peak at ~7.5 min corresponds to the expected product. The peak at ~6.6 min is present in the control without substrate. LC-MS method 2 was used for this analysis.

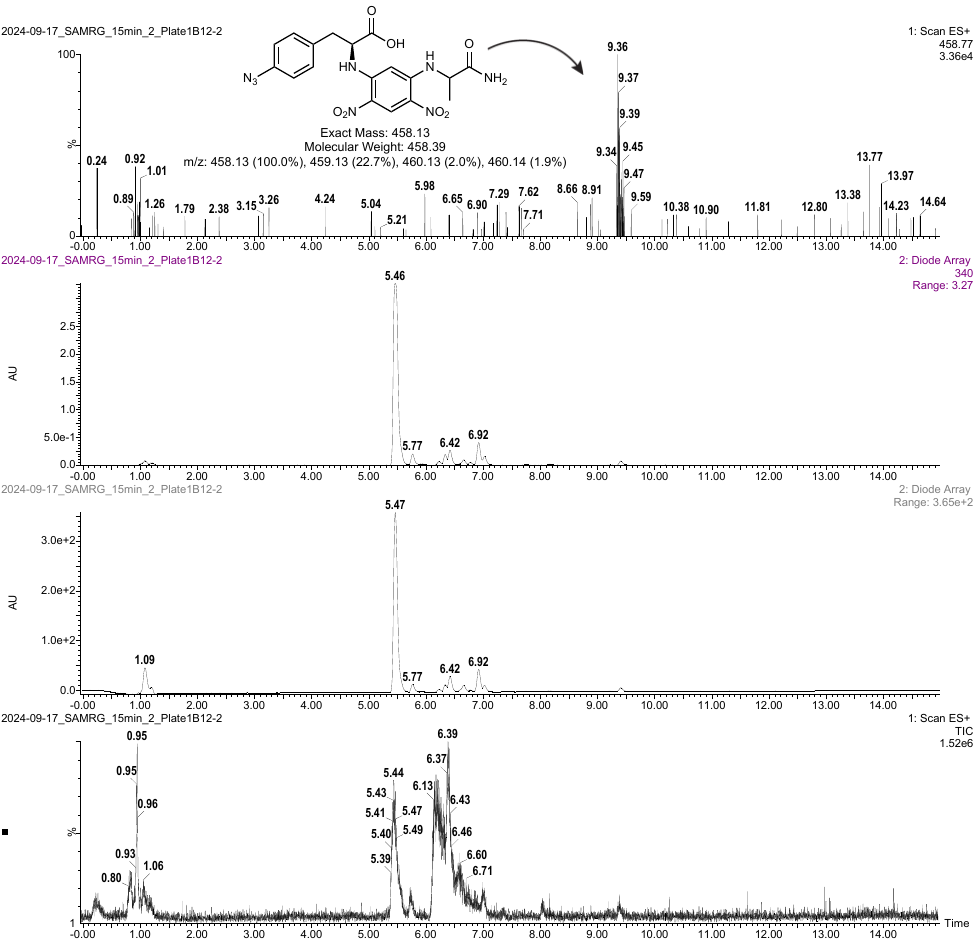

Figure S7. Marfey’s analysis on the one-pot s-ObiH, *Rp*PSDH, TyrB catalyzed reaction with **4b**. LC-MS method 2 was used for this analysis.

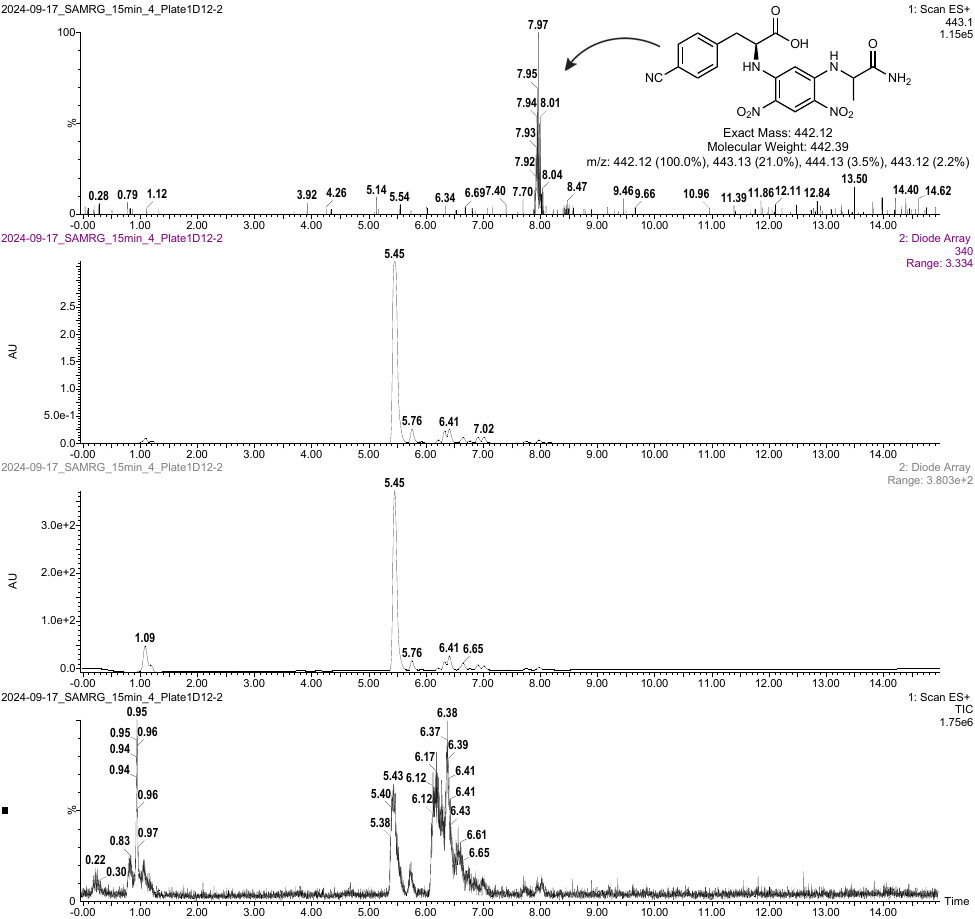

Figure S8. Marfey’s analysis on the one-pot s-ObiH, *Rp*PSDH, TyrB catalyzed reaction with **5b**. LC-MS method 2 was used for this analysis.

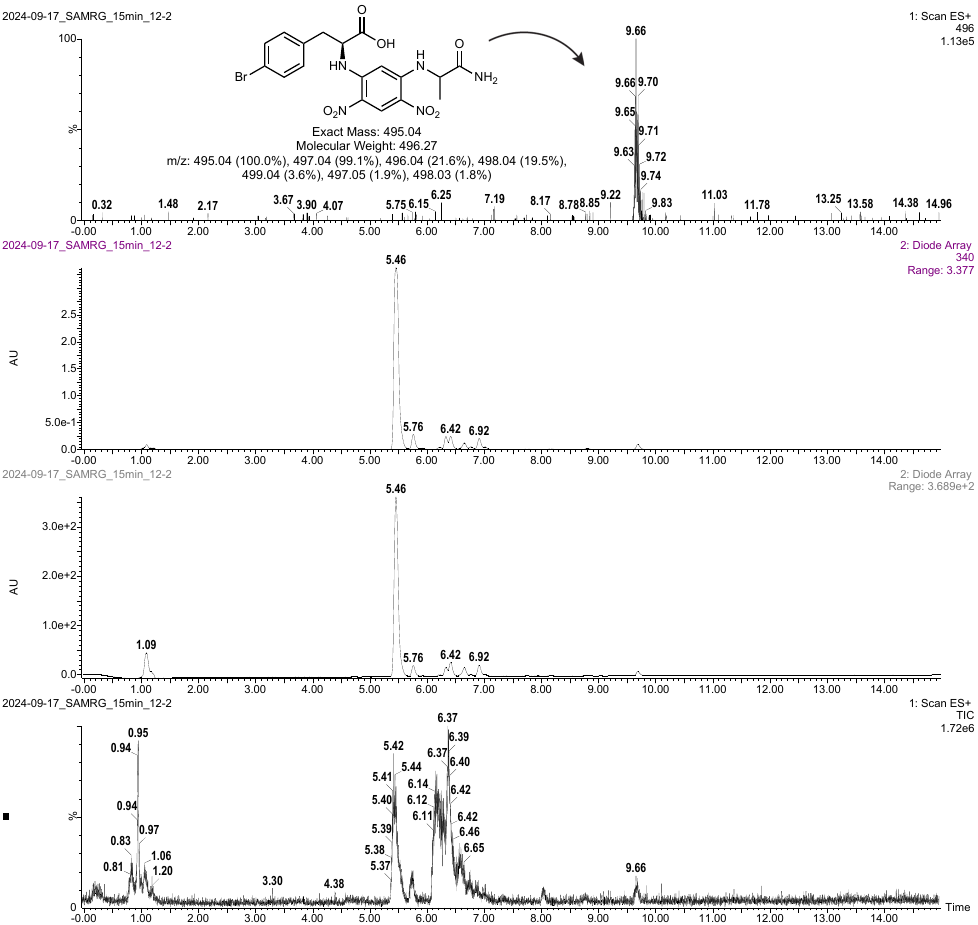

Figure S9. Marfey’s analysis on the one-pot s-ObiH, *Rp*PSDH, TyrB catalyzed reaction with **6b**. LC-MS method 2 was used for this analysis.

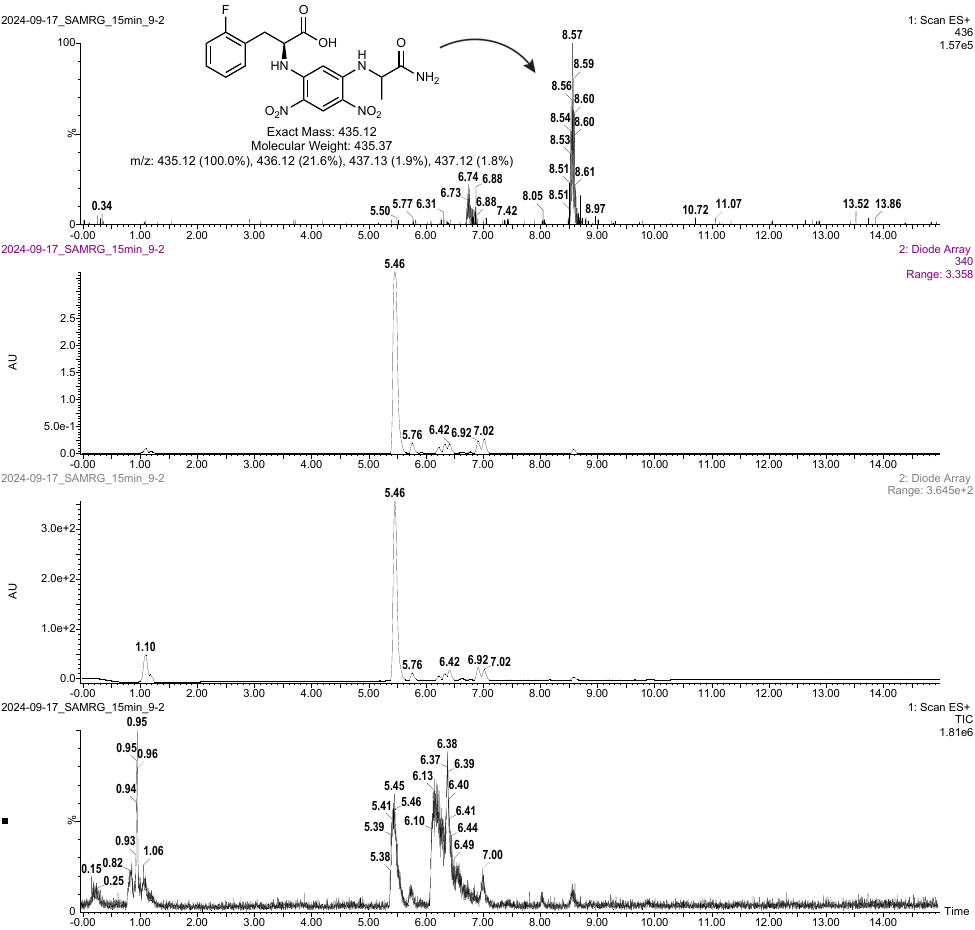

Figure S10. Marfey’s analysis on the one-pot s-ObiH, *Rp*PSDH, TyrB catalyzed reaction with **7b**. LC-MS method 2 was used for this analysis.

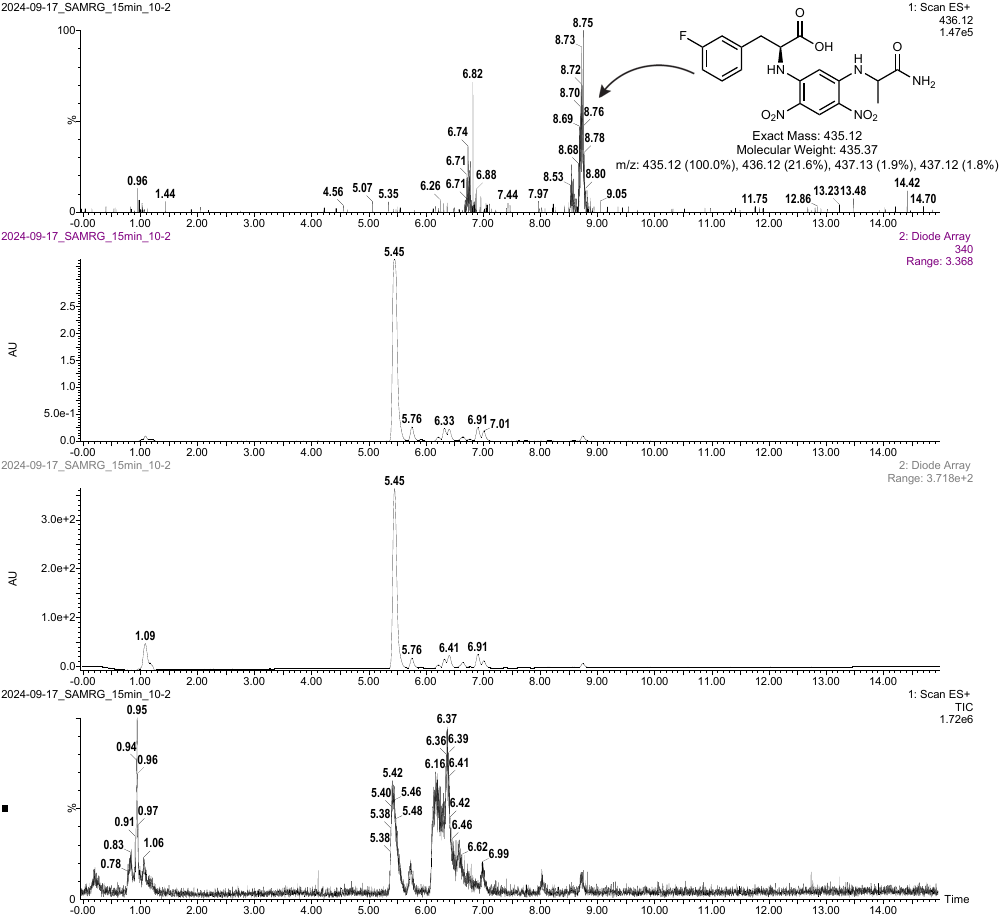

Figure S11. Marfey’s analysis on the one-pot s-ObiH, *Rp*PSDH, TyrB catalyzed reaction with **8b**. LC-MS method 2 was used for this analysis.

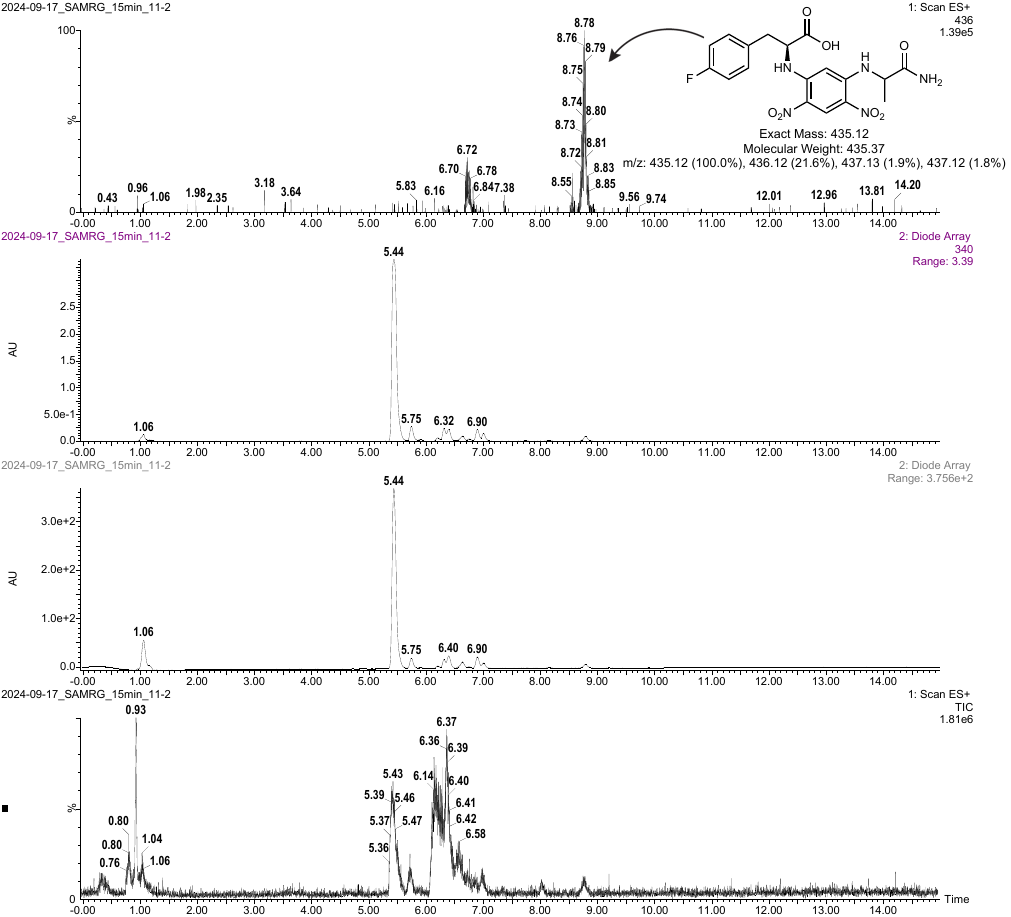

Figure S12. Marfey’s analysis on the one-pot s-ObiH, *Rp*PSDH, TyrB catalyzed reaction with **9b**. LC-MS method 2 was used for this analysis.

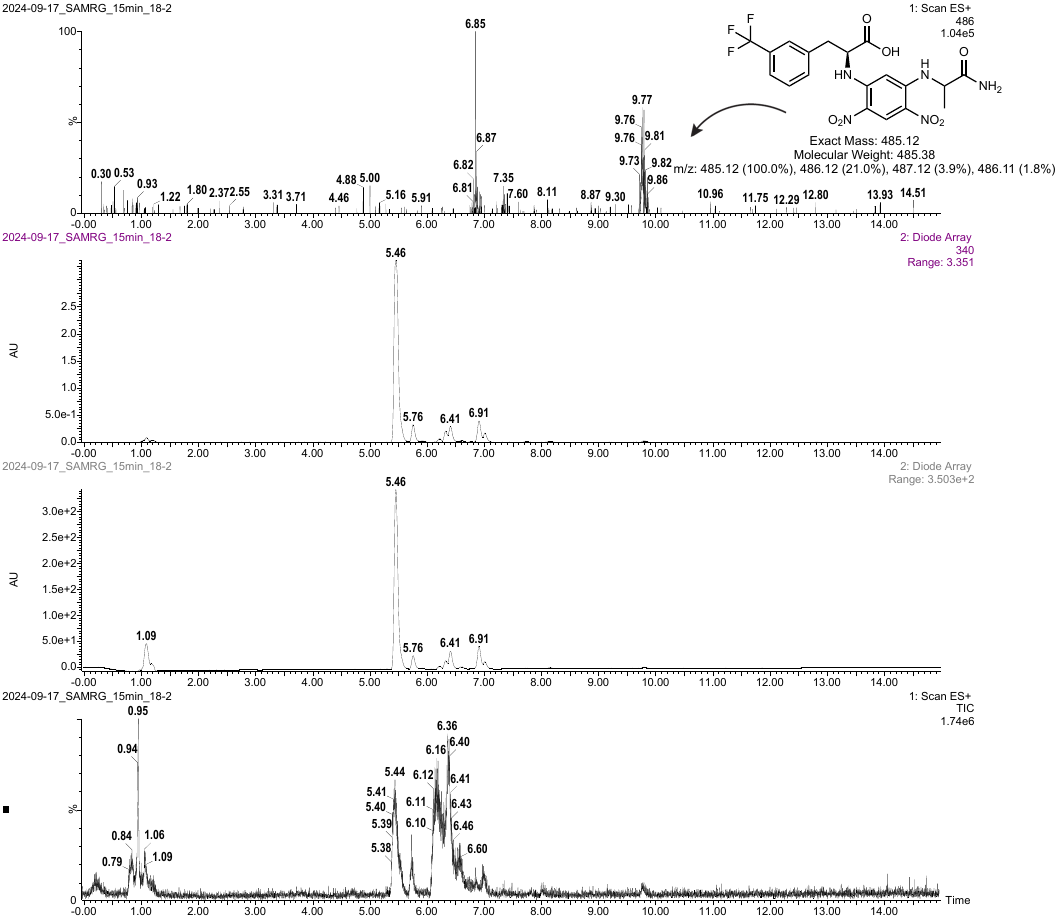

Figure S13. Marfey’s analysis on the one-pot s-ObiH, *Rp*PSDH, TyrB catalyzed reaction with **10b**. LC-MS method 2 was used for this analysis.

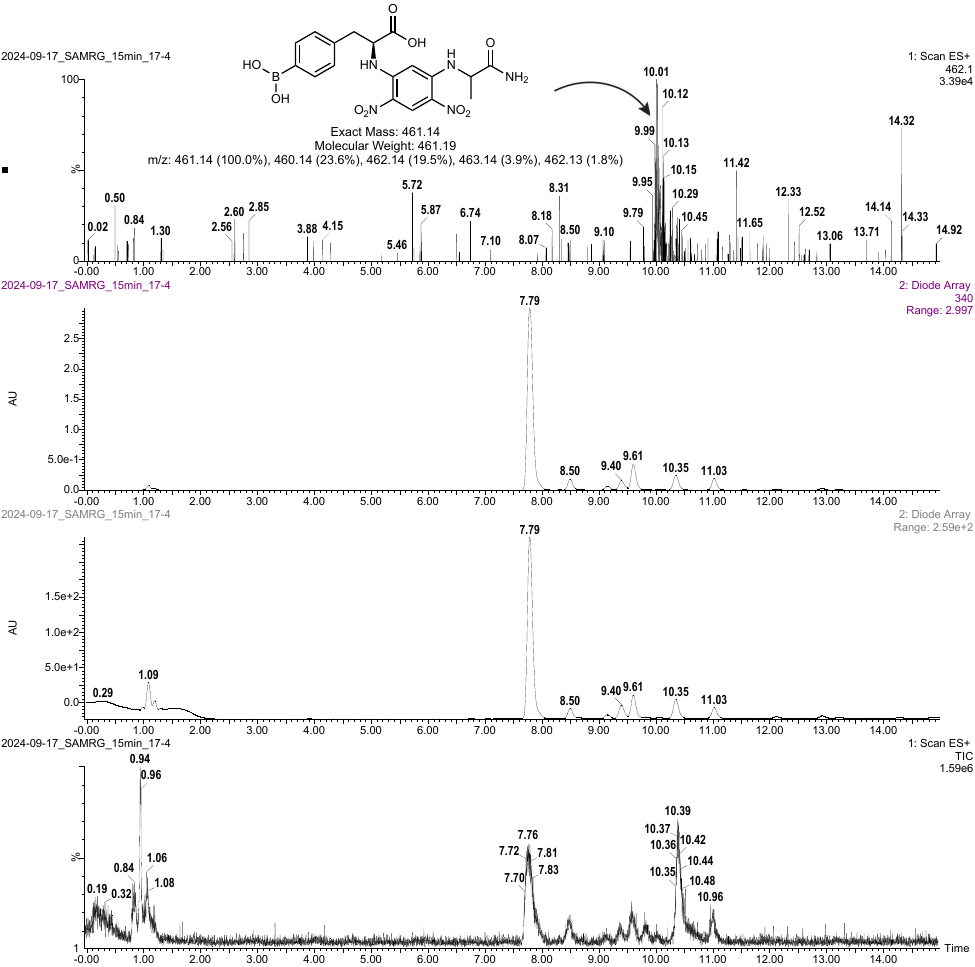

Figure S14. Marfey’s analysis on the one-pot s-ObiH, *Rp*PSDH, TyrB catalyzed reaction with **12b**. LC-MS method 3 was used for this analysis.

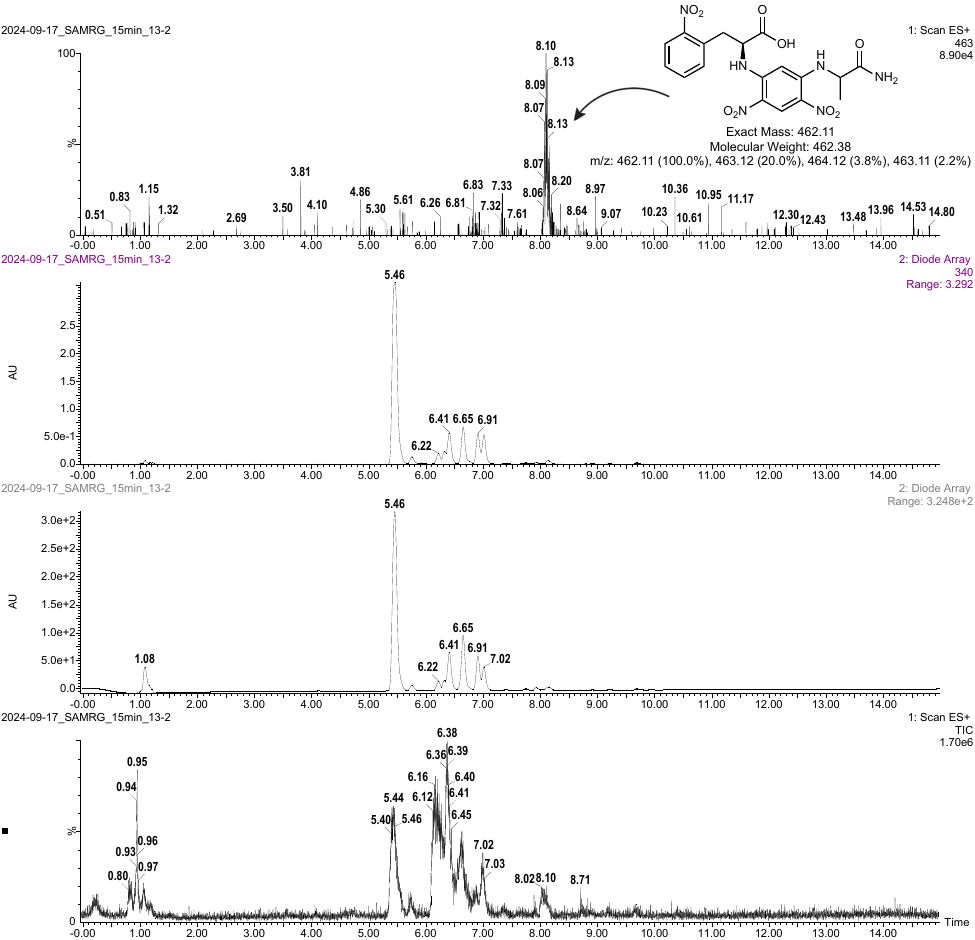

Figure S15. Marfey’s analysis on the one-pot s-ObiH, *Rp*PSDH, TyrB catalyzed reaction with **13b**. LC-MS method 2 was used for this analysis.

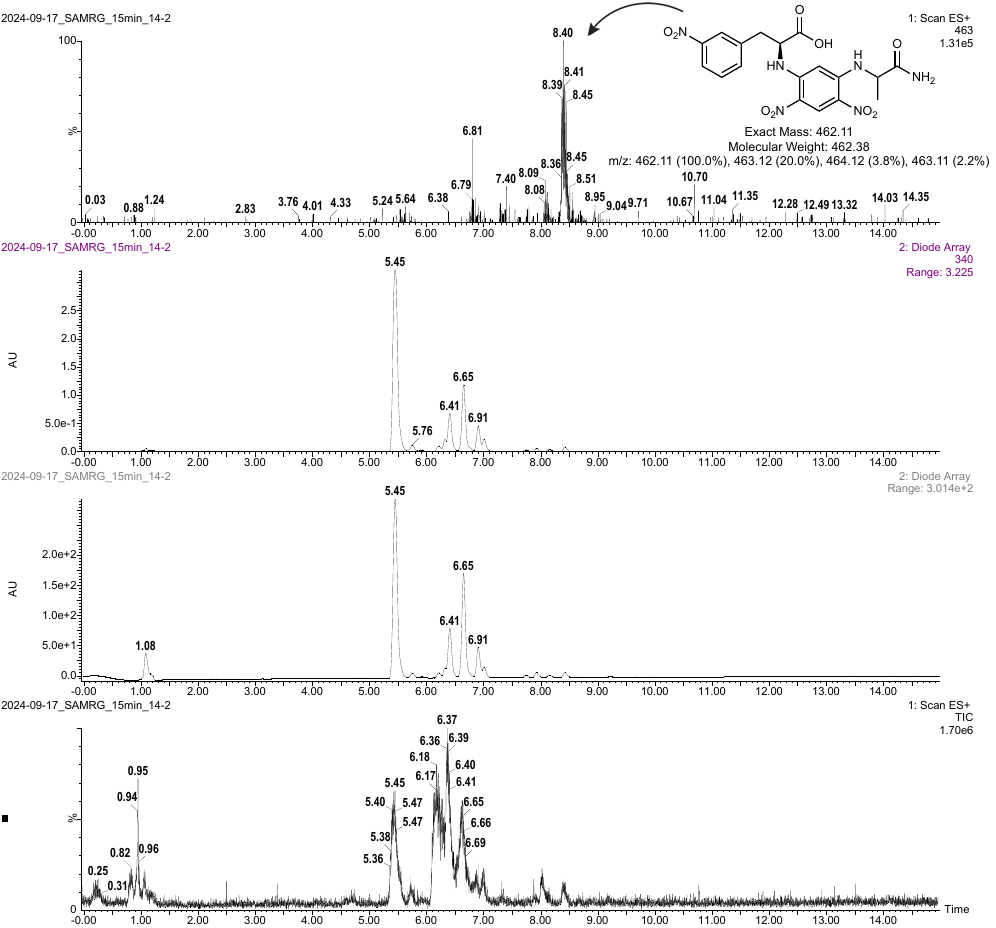

Figure S16. Marfey’s analysis on the one-pot s-ObiH, *Rp*PSDH, TyrB catalyzed reaction with **14b**. LC-MS method 2 was used for this analysis.

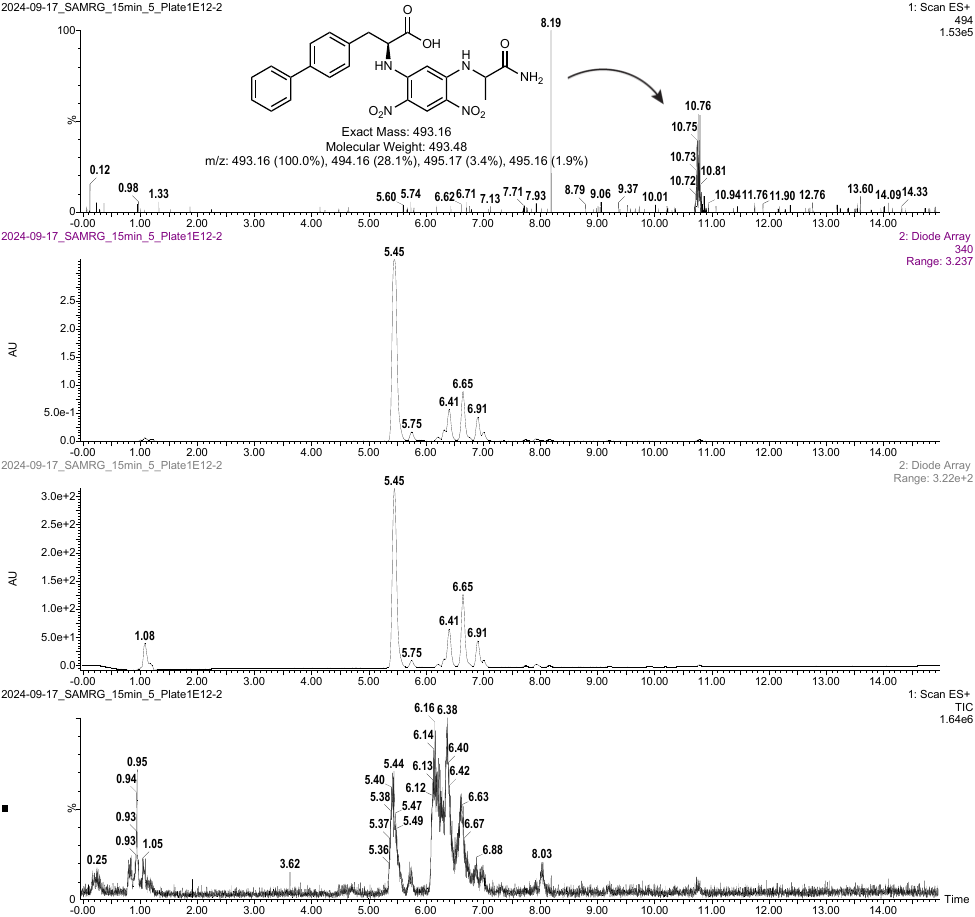

Figure S17. Marfey’s analysis on the one-pot s-ObiH, *Rp*PSDH, TyrB catalyzed reaction with **15b**. LC-MS method 2 was used for this analysis.

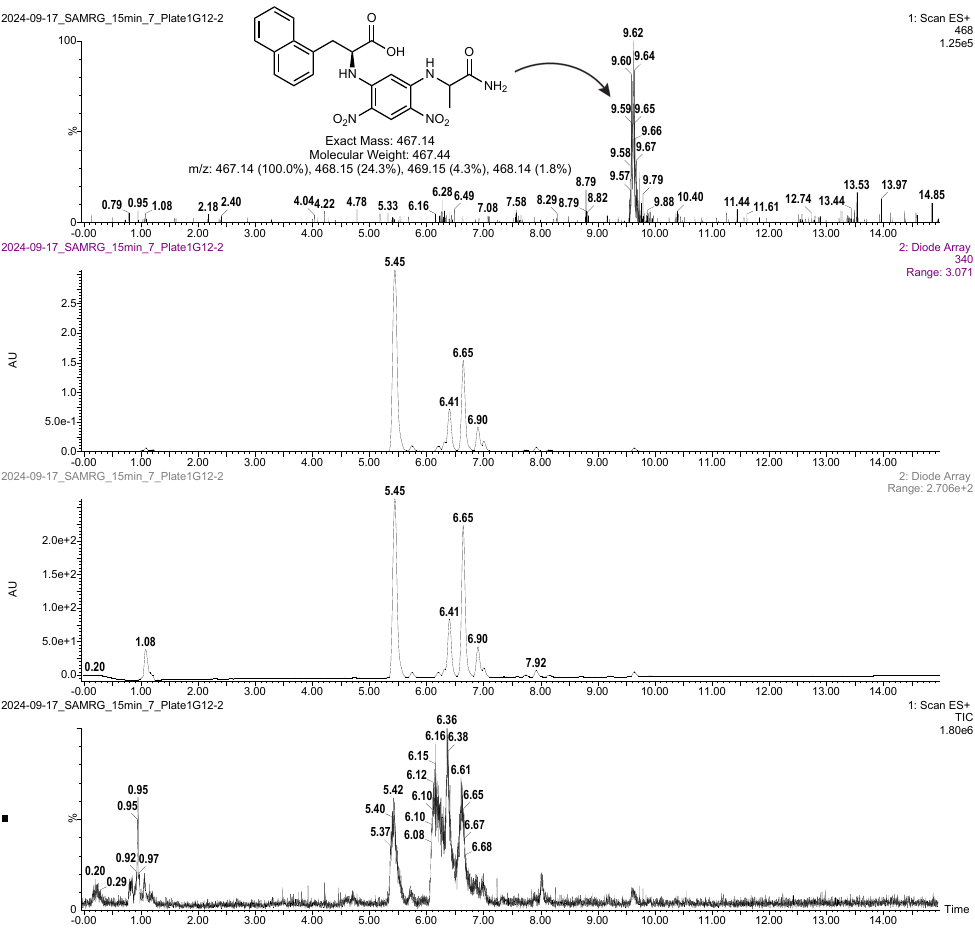

Figure S18. Marfey’s analysis on the one-pot s-ObiH, *Rp*PSDH, TyrB catalyzed reaction with **16b**. LC-MS method 2 was used for this analysis.

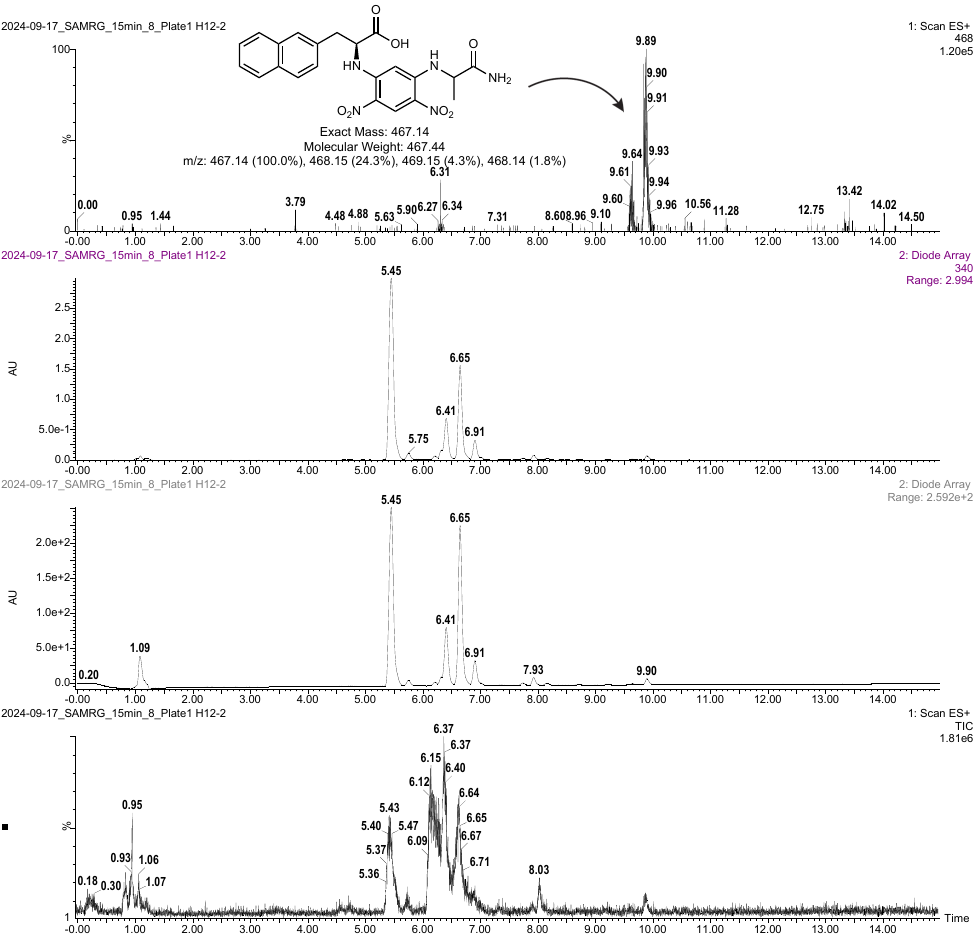

Figure S19. Marfey’s analysis on the one-pot s-ObiH, *Rp*PSDH, TyrB catalyzed reaction with **17b**. LC-MS method 2 was used for this analysis.

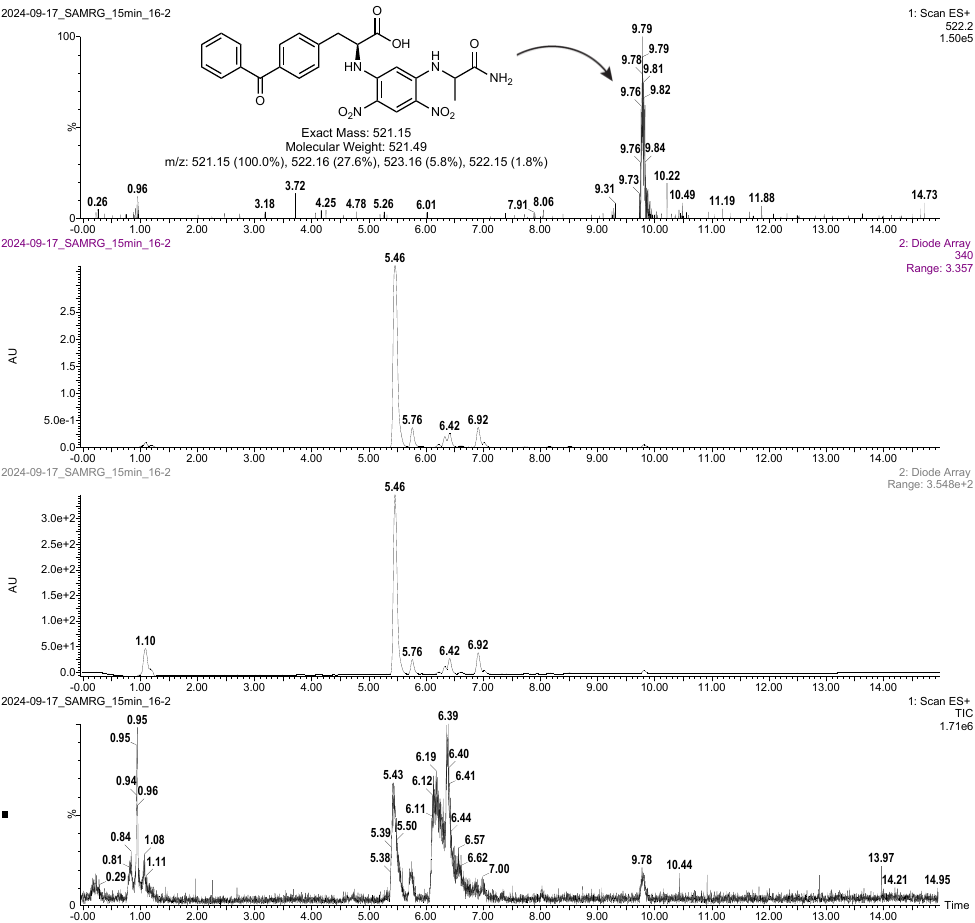

Figure S20. Marfey’s analysis on the one-pot s-ObiH, *Rp*PSDH, TyrB catalyzed reaction with **18b**. LC-MS method 2 was used for this analysis.

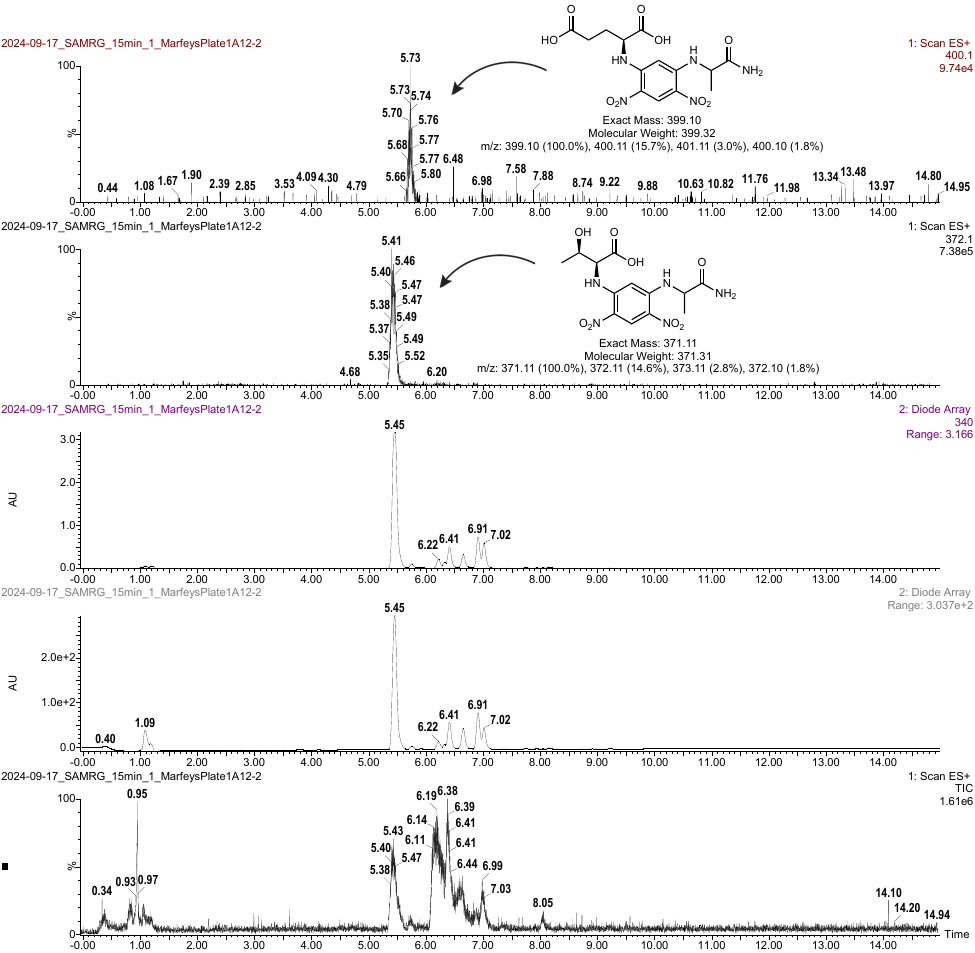

Figure S21. Marfey’s analysis on the one-pot s-ObiH, *Rp*PSDH, TyrB reaction with no substrate (control). The derivatized L-Glu and L-Thr in the reaction mixture are shown. LC-MS method 2 was used for this analysis.

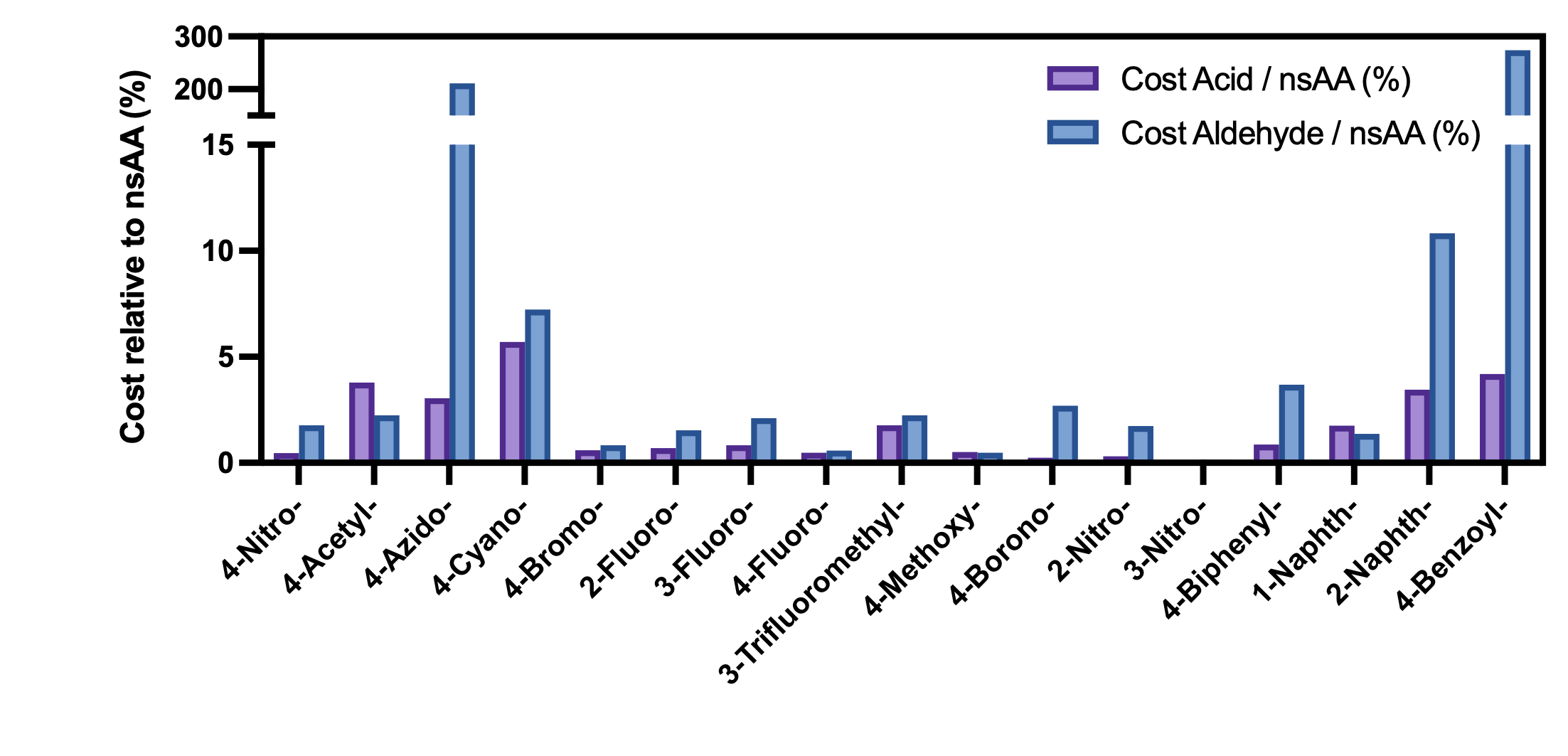

Figure S22. Cost analysis of the phenylalanine derivative precursors (aldehydes or acids) relative to the nsAA product. The cost of each chemical was determined on a cost per mol basis. Prices were compiled in Q1 2024 from the following chemical vendors: Thermo Fisher Scientific, Santa Cruz Biotechnology, AA Blocks, and Ambeed. Most aldehydes (15 out of the 17 analyzed) were less than 11% of the cost relative to the corresponding nsAA analogs. The exceptions in which the aldehyde was more costly than the nsAA include 4-azidobenzaldehyde and 4-benzoylbenzaldehyde. All acids analyzed were less than 6% of the cost relative to the corresponding nsAA analogs

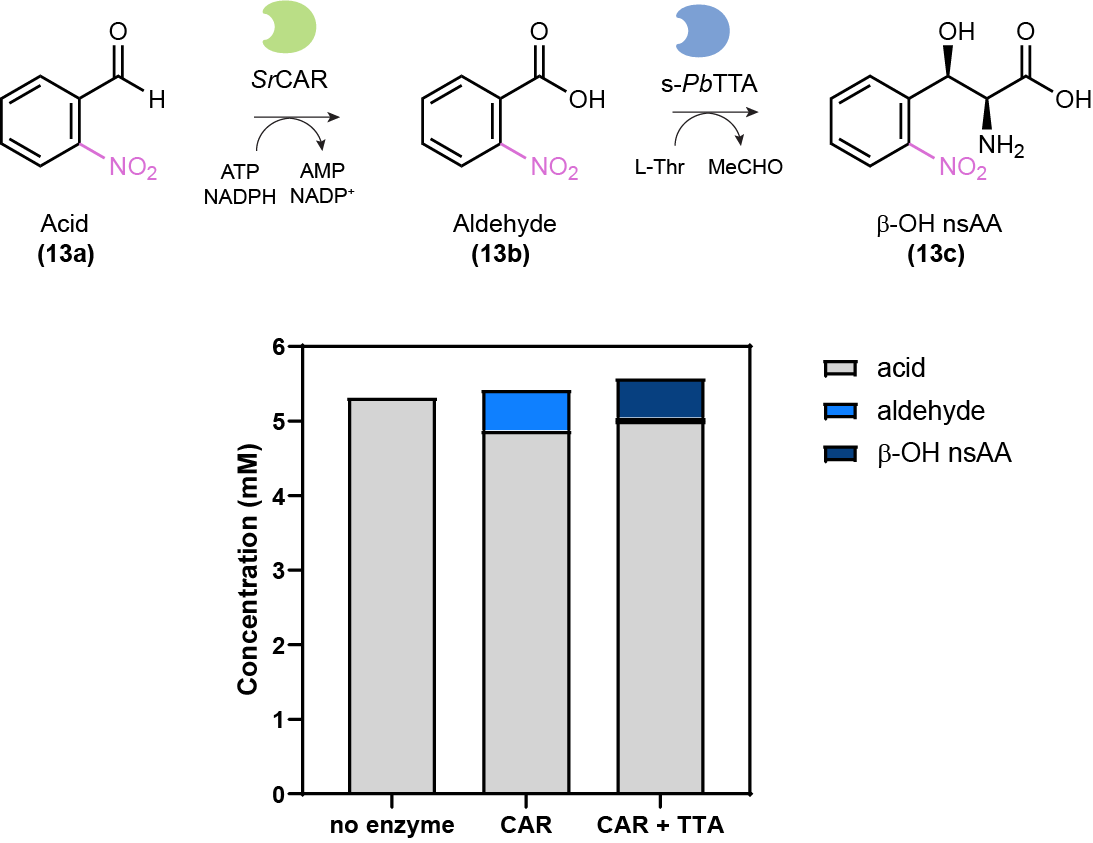

Figure S23. Endpoint conversion of 5 mM 2-nitrobenzoic acid (**13a**) supplied to the early steps of the biocatalytic cascade. The reaction was performed with 4 µM SrCAR (0.52 mg/mL), 1 µM s-PbTTA (0.06 mg/mL), 100 mM HEPES pH 8, 400 µM PLP, 20 mM MgCl_2_, 1.5 mM ATP, 1.5 mM NADPH, 100 mM L-Thr, 5% (v/v) DMSO, and 5 mM of acid tested (100 mM stock in DMSO) for 23 h at 30 ºC at 1000 RPM. When only CAR is provided, the anticipated product would be 2-nitrobenzaldehyde (**13b**). When both CAR and L-TTA are combined, the anticipated product would be the corresponding beta-hydroxylated alpha-amino acid (β-OH nsAA). HPLC measurement of the concentration of carboxylic acid, aldehyde, or β-OH nsAA shows that unreacted 2-nitro-benzoic acid is the dominant component, indicating that low *Sr*CAR activity on this substrate (**13a**) is responsible for the low 2-nitro-phenylalanine yield observed in Figure 2. The L-TTA used for this experiment was a N-terminal His_6_ and SUMO-tagged variant from *Parachlamydiales bacterium* (s-*Pb*TTA). Synthesis of the chemical standard for the β-OH nsAA is detailed in Jones, *et al*.

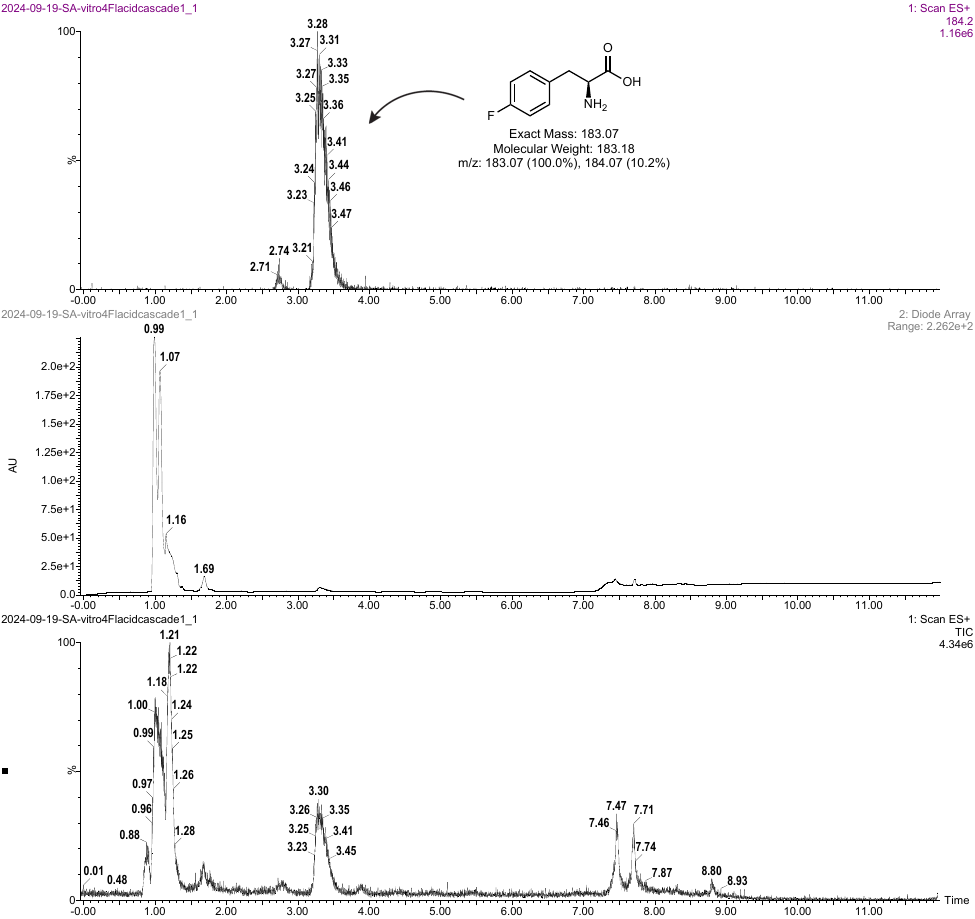

Figure S24. Production of 4-fluoro-phenylalanine (**9e**) from the 4-fluoro-benzoic acid (**9a**) precursor, as confirmed by MS.

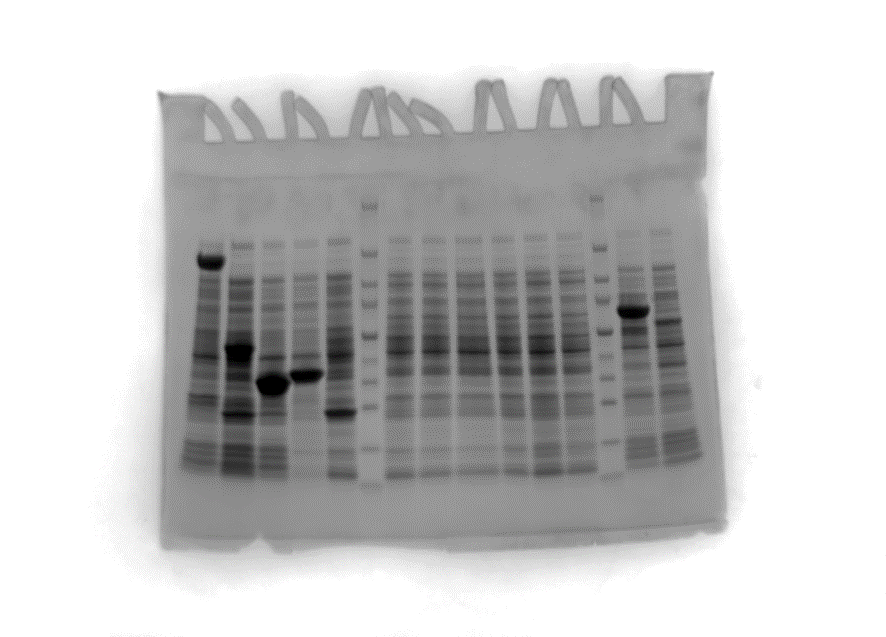

Figure S25. SDS-PAGE of clarified lysates. Left = ladder (Biorad Precision Plus Unstained Protein Standard (Catalog # 1610363)), middle = lysate prepared from the RARE.∆16 strain expressing s-ObiH (62.6 kDa), right = lysate prepared from the RARE.∆16 strain expressing s-*Rp*PSDH (48.6 kDa).

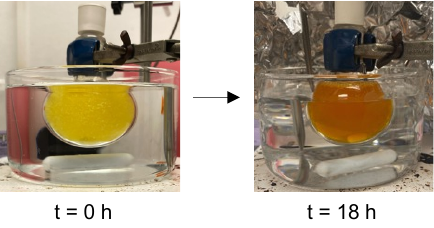

Figure S26. Preparative-scale nsAA synthesis using clarified lysate. By supplementing 25 mM of terephthaldehyde (**3b**) to a 40 mL reaction containing s-ObiH and s-*Rp*PSDH clarified lysates (reaction shown in Figure 3d), we observed a progressive color change from light yellow to a deeper orange as the reaction proceeded over 18 h.

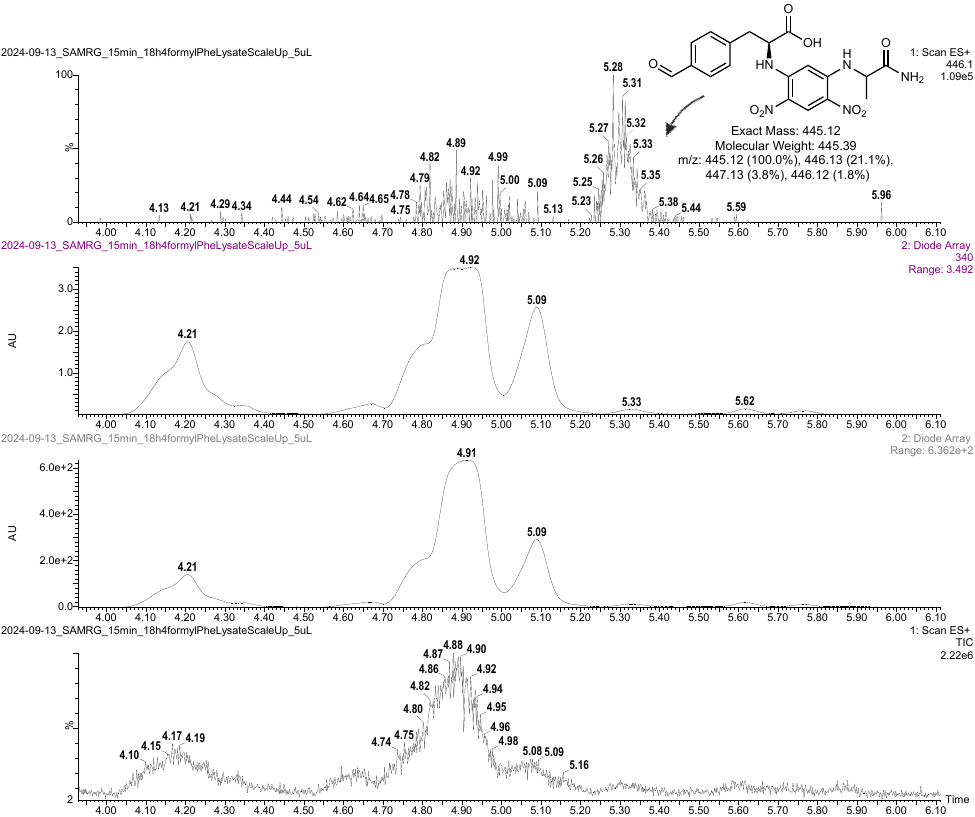

Figure S27. Marfey’s analysis on the one-pot s-ObiH lysate, s-*Rp*PSDH lysate catalyzed reaction with **3b**. LC-MS method 1 was used for this analysis.
